# Distinct medial-temporal lobe mechanisms of encoding and amygdala-mediated memory reinstatement for disgust and fear

**DOI:** 10.1101/825844

**Authors:** Monika Riegel, Małgorzata Wierzba, Marek Wypych, Maureen Ritchey, Katarzyna Jednoróg, Anna Grabowska, Patrik Vuilleumier, Artur Marchewka

## Abstract

Remembering events that evoke emotions such as disgust or fear is critical to our survival. However, previous studies investigating the interplay between emotion and memory disregarded the effects of specific emotions, leading to inconsistent results. Also, the role of amygdala throughout memory stages has been poorly understood. Here, we show that after 3 weeks delay, word pairs evoking disgust were remembered better than pairs evoking fear. These two emotions distinctly modulated neural mechanisms of memory. Successful encoding of disgust-evoking information was mediated by univariate activation in amygdala and perirhinal cortex, in contrast to fear-evoking memories that engaged hippocampus and parahippocampal gyrus. Critically, univariate activation in the amygdala during encoding was correlated with memory reinstatement of individual word pairs, and more so for disgust than for fear. Together, these findings shed a new light on the role of the amygdala and medial temporal lobe regions in encoding and reinstatement of specific emotional memories.

## Introduction

Although unpleasant, experiencing emotions such as disgust and fear is crucial to our lives. Both are negative, arousing and motivating avoidance, and their primary function is protecting us from threat or contamination. For instance, getting assaulted in a park around the block would evoke your fear, whereas finding a cockroach in your favourite cereals would evoke your disgust. As a result, you would be motivated to avoid the park or the cereals in the future. To make it happen, you need to form durable memories of complex events evoking these essential emotions.

However, the previous studies have shown inhomogenous effects of emotion on memory for associations between the pieces of information, including either *enhancement, impairment*, or *null effects* (reviews: Chiu, Dolcos, Gonsalves, & Cohen, 2013; Murray & Kensinger, 2013; Yonelinas & Ritchey, 2015). This inconsistency might depend on at least two factors, namely how close are the associations formed between items and what is the type of emotion. At the same time, the role of amygdala – a critical hub in emotion processing – was studied selectively during encoding or retrieval. However, memory requires interactions between these two memory stages.

First, when complex between-item associations have to be formed, emotion was reported to *impair* memory (Bergmann, Rijpkema, Fernández, & Kessels, 2012; Murray & Kensinger, 2012; Pierce & Kensinger, 2011; Rimmele et al., 2011). However, emotion may *enhance* associative memory when the information is merged and processed as an within-item association, or even as one single item, reflecting so-called unitization (Graf & Schacter, 1989; Murray & Kensinger, 2013). This continuous process is influenced by characteristics of the to-be-merged items and the encoding task. For instance, encoding instructions requiring mental integration trigger active unitization attempts more so than non-integrative encoding instructions (Madan, Fujiwara, Caplan, & Sommer, 2017; Murray & Kensinger, 2014a).

Second, to date, memory modulation by emotion was explained mostly in relation to arousal and valence, even though behavioral differences were observed for distinct emotion categories, such as disgust and fear (Chapman, Johannes, Poppenk, Moscovitch, & Anderson, 2012; Croucher, Calder, Ramponi, Barnard, & Murphy, 2011; van Hooff, Devue, Vieweg, & Theeuwes, 2013; van Hooff, van Buuringen, El M’rabet, de Gier, & van Zalingen, 2014). Both are negative, arousing and motivating avoidance (Krusemark & Li, 2011; Woody, 2000). At the same time, they induce distinct physiological responses (Cisler, Olatunji, & Lohr, 2009; Critchley et al., 2005) and have different effects on cognitive processes. Emotion category-specific effects on memory for associations has not been investigated so far, but a better recall of disgust-compared to fear-related stimuli was reported for single words (Charash & McKay, 2002) and images (Croucher et al., 2011). These differences might reflect dissociable components implicated in the memory for different emotion categories (Berridge, 2019), perhaps evolutionary-based(Susskind et al., 2008), and possibly supported by distinct brain mechanisms.

Modulation of emotional memories is known to involve the amygdala (AMY) via its interaction with other medial temporal lobe (MTL) regions (Cahill & McGaugh, 1998; Kensinger & Schacter, 2006). Animal research has shown that the AMY shares strong reciprocal connections with anterior hippocampus (HC), parahippocampal cortex (PHC) and perirhinal cortex (PRC) (Pitkänen, Pikkarainen, Nurminen, & Ylinen, 2000), each having a different function in memory formation. Along with PRC receiving projections from the ventral visual ‘what’ stream (processing of items and objects in the environment), PHC receives projections from the dorsal ‘where’ stream (processing of contextual information, for instance spatial and temporal) (Ranganath, 2010), whereas HC binds item and context information that make up an event (Ranganath, 2010). For neutral information, encoding and retrieval of unitized compared to non-unitized pairs was shown to rely less on HC binding, and more on familiarity-based processes mediated by PRC (Haskins, Yonelinas, Quamme, & Ranganath, 2008; Memel & Ryan, 2017, 2018).

But how this functional specialization within the MTL could be influenced by emotion modulation and AMY engagement? Studies have demonstrated an increase in functional connectivity between right HC and AMY during encoding of negatively valenced associations of faces and identities (Berkers, Klumpers, & Fernández, 2015) when compared to neutral ones. On the other hand, the arousal effect on memorizing unitized words was shown in negative correlation between the activation of left AMY, frontal areas, and HC (Murray & Kensinger, 2014a). Finally, most recent study provided evidence for a double dissociation, with AMY and PRC activity during encoding supporting the recollection advantage for single emotional items, whereas HC and PHC activity predicting subsequent context memory for both emotional and neutral pictures (Ritchey, Wang, Yonelinas, & Ranganath, 2019). However, the effects of emotion and AMY engagement on memory for associations was only explained in terms of arousal and valence, while differences between basic emotion categories were generally disregarded. Moreover, it was mostly investigated during encoding, whereas models of episodic memory suggest that remembering involves the reenactment of encoding processes during retrieval (Danker & Anderson, 2010). How emotion and AMY engagement determines subsequent reinstatement has been unknown.

Here we determined whether the emotional memory enhancement occurs over long-term delays for unitized word pairs, and if so, whether it is differentially sensitive to distinct emotion categories (disgust and fear) irrespective of their valence and arousal levels. We investigated brain mechanisms hypothesized to occur within MTL regions recruited during encoding and reinstatement, for both disgust- and fear-related unitizations. To do so, we combined a recognition memory task (Radvansky, 2006) with functional magnetic resonance (fMRI) and three analytical approaches: univariate activation analysis, functional connectivity analysis and representational similarity analysis (RSA, Kriegeskorte, 2008). During encoding, participants mentally unitized and memorized semantically congruent Polish word pairs (35 per condition) associated with disgust (e.g. *swab – stinky*), fear (e.g. *throat – knife*), or emotionally neutral (e.g. *circle – axis*). After 2-3 weeks, they performed a recognition test during which novel word pairs were also presented as lures (25 per condition). Both phases took place during fMRI scanning. Critically, in a final phase of the experiment, participants rated emotions evoked by the word pairs, so that we could individually control for the effects of valence and arousal dimensions. Most importantly, this paradigm enabled us to compare the encoding-retrieval similarity of brain activation patterns elicited across two different emotion categories and determine which brain regions promoted this measure of neural reinstatement.

We found a memory enhancement effect due to negative emotion. Moreover, word pairs related to disgust were remembered better than word pairs related to fear, and this effect could not be explained by differences in arousal. Furthermore, we found distinct brain pathways preferentially engaged in memory modulation by disgust or fear. Specifically, the left AMY and left PRC were more activated during successful encoding of disgust-related unitizations, whereas right PHC and HC were more activated for fear-related unitizations. As previously, these differences are observed over and above the effects of arousal. Moreover, during successful encoding of disgust-compared to fear-related word pairs, we observed an increase in functional connectivity between AMY, PRC and PHC. Finally, the left AMY encoding activation was correlated with the HC activation during retrieval, as well as with memory reinstatement of emotion-specific representations of particular word pairs. This correlation was higher for disgust than fear.

Altogether, the results show that behavioural and neuronal mechanisms of long-term memory for associations are differently modulated by distinct emotion categories, over and above valence and arousal dimensions. For the first time, we revealed that due to AMY and PRC engagement, disgust as opposed to fear has a strong influence on encoding and memory reinstatement of verbal information. This finding is in line with its distinctive and perhaps evolutionary-driven role in memorizing and avoiding possible sources of contamination. The current results provide an important complement to debates on theoretical frameworks for emotion and cognition research by showing basic emotion effects beyond affective dimensions.

## Results

### Emotion-specific effects on memory

Recognition memory was tested 15-19 days (M = 16.86, SD = 1.22) after encoding. Among old word pairs, both emotional categories were correctly recognized more often than neutral word pairs, and word pairs related to disgust were correctly recognized more often than those related to fear (Supplementary Fig. S1). To take into account possible responding strategies of the participants, we also computed the sensitivity index (d’) derived from signal detection theory (SDT) and confirmed significant differences in d’ between emotion categories [F(2,102) = 20.20, p < .001, η^2^= .28]. Disgust-related word pairs were recognized with greater sensitivity than fear-related (p < .001) and neutral pairs (p < .001). There was no significant difference in d’ between fear-related and neutral pairs (Fig. 1a).

**Fig. 1.**
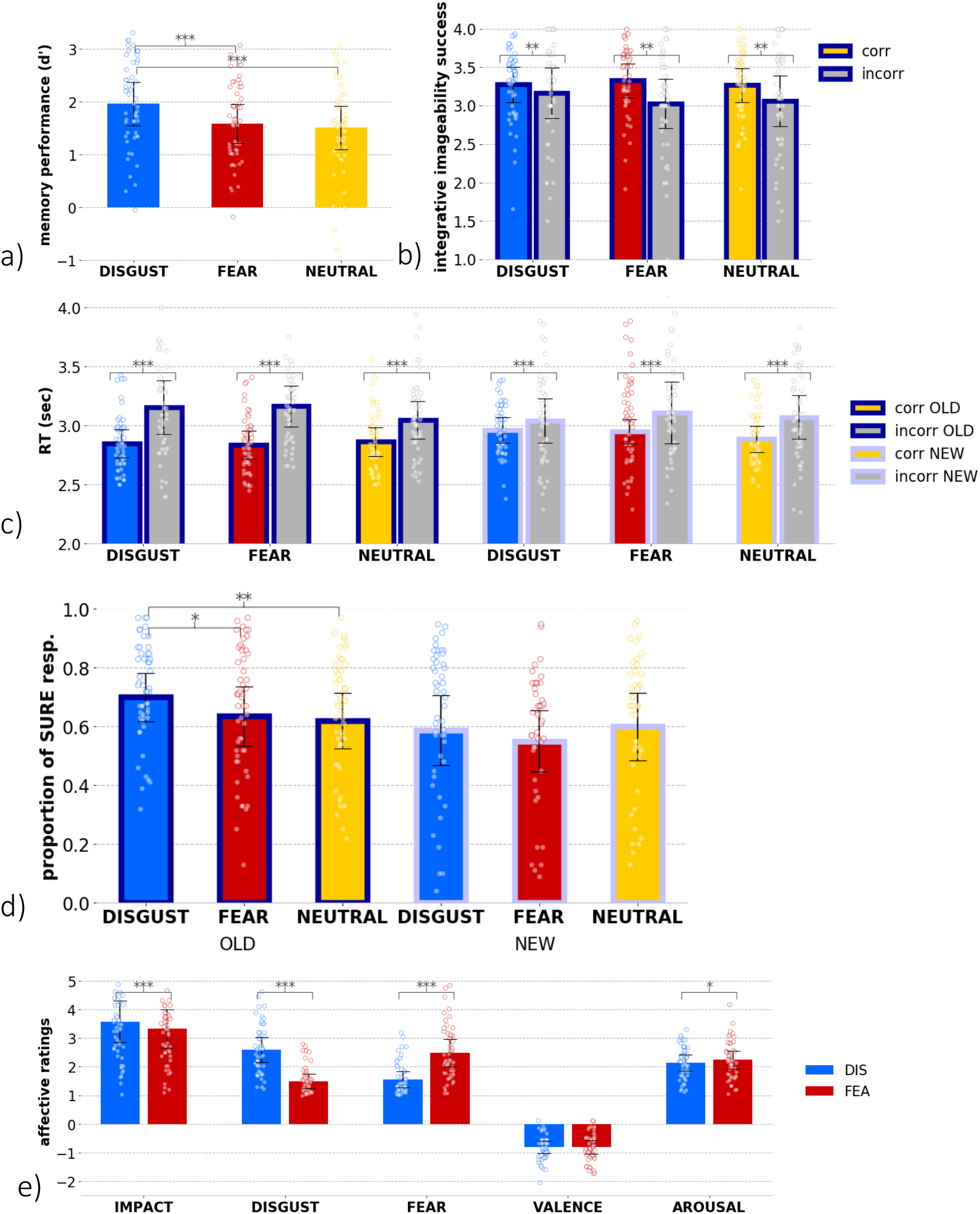
a) Memory performance (sensitivity index, d’) obtained for word pairs representing each emotion category; b) Integrative imageability success obtained for correctly and incorrectly recognized old word pairs representing each emotion category; c) Reaction times (RT) in the recognition task obtained for correctly and incorrectly recognized old and new word-pairs representing each emotion category; measured in seconds from the word pair’s onset until a response; d) Sureness of recognition obtained for correctly recognized old and new word pairs representing each emotion category; expressed in proportion (%) of SURE responses among all responses; e) Affective ratings of impact, disgust, fear, valence, and arousal obtained for disgust- and fear-related word pairs; error bars represent one standard deviation, dots represent individual subjects’ scores; corr – correct, incorr – incorrect, DIS – disgust, FEA – fear; * p < .05, ** p < .005, *** p < .001.

**Fig 2.**
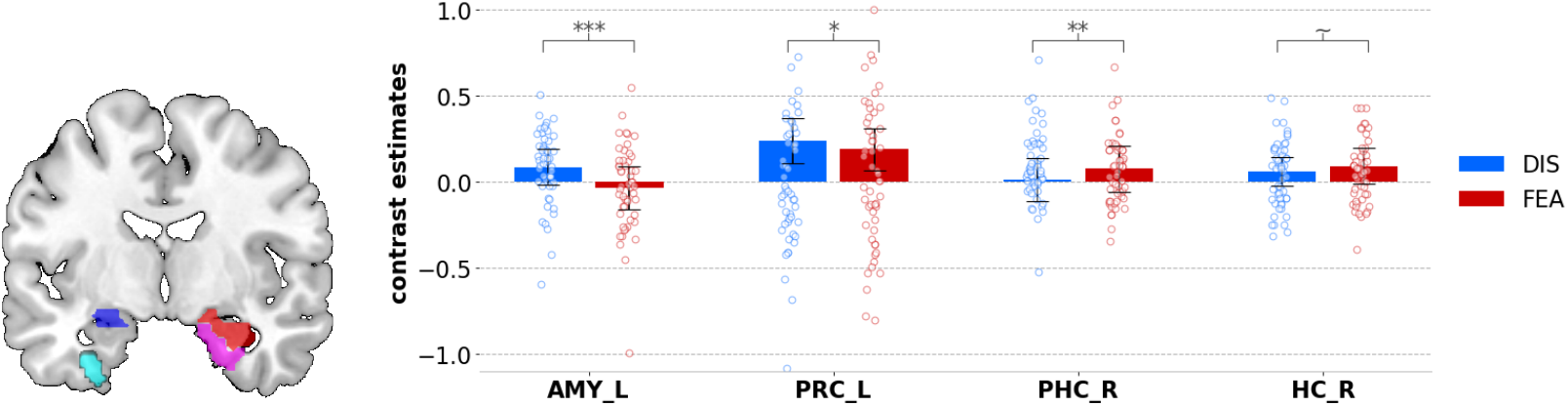
Emotion-related differences in encoding activity. Bars represent contrast estimates for successful encoding activity observed in presented anatomical ROIs (marked on the brain template: left AMY in blue, left PRC in cyan; right PHC in violet and right HC in red), split according to emotion factor (disgust- and fear-related word pairs); error bars represent one standard deviation, dots represent individual subjects’ scores; DIS – disgust, FEA – fear; ~ trend toward significance, * p < .05, ** p < .005, *** p < .001.

To confirm emotion category-specific effects (i.e. better recognition of disgust-vs. fear-related word pairs), we controlled for the level of arousal associated with each condition by including the difference in arousal ratings (see: Fig. 1e) for all word pairs as a covariate in the analysis described above. The results still showed the main effect of emotion [F(1,50) = 19.79, p < .001, η^2^ = .28] with higher sensitivity index (d’) for disgust than fear (p < .001). This finding emphasizes that differences in memory performance observed for these two basic emotion categories were not driven by differences in emotional arousal.

Reaction times (RT) during recognition were also analyzed to control for the possible differences in task difficulty among emotion conditions. Only interaction between correctness and novelty (old, new) of word pairs was found [F(1,45) = 3.97, p = .05, η^2^= .08], such that correct recognition of old word pairs was significantly faster than new word pairs, but there was no difference in the case of incorrect recognition, as depicted in Fig. 1c. These effects were independent of emotion categories (disgust, fear, neutral) and we can infer that recognition task was not harder for any emotion category.

To determine whether emotion effects on memory could be related to the strength of associations created between words, we then analyzed the ratings of integrative imageability success given by the participants during encoding. The results showed that correctly recognized word pairs were better integrated during encoding compared to those recognized incorrectly [F(1,45) = 11.086, p = .002, η^2^ = .198], again regardless of emotion category (disgust, fear, neutral). Thus, the differential memory modulation by emotion categories did not result from the strength of associations created during encoding (Fig. 1b).

Next, we analyzed how the memory and emotion effects were related to subjective confidence, or sureness, rated by the participants for each correct recognition response. We found that the proportion of sure responses was significantly modulated by emotion category (disgust, fear, neutral) and by novelty (old, new) of the word pairs (interaction effect [F(2,78) = 4.309, p = .017, η^2^ = .099]), with more frequent sure responses to disgust-than fear-related (p = .013) and neutral word pairs (p = .003) among old pairs (Fig 4). Among new word pairs, no significant differences were found (Fig. 1d). In general, the emotion and novelty effects on subjective confidence of recognition showed a similar pattern to their effect on the recognition rate – the better remembered, the higher confidence.

Finally, affective ratings collected from the participants were analyzed to determine any potential differences between word pairs assigned to different emotion categories (disgust, fear, neutral). As expected, we found that all the affective parameters were higher (p < .001) for both disgust- and fear-related word pairs than for neutral pairs. The ratings of disgust were significantly higher for disgust-related than fear-related (t = 12.53, p < .001) pairs, whereas ratings of fear were lower for disgust-than fear-related (t = −12.42, p < .001) pairs, which demonstrates the validity of our experimental manipulation. There was no significant difference in valence ratings between them. Unexpectedly, however, significant differences in the arousal and impact (for definition see: Methods) ratings were found. Specifically, the level of arousal was rated as lower for disgust-related than fear-related (t = −2.73, p = .009) word pairs, whereas impact was rated higher for disgust-than fear-related (t = 3.78, p < .001) pairs (Fig. 1e). Therefore, these differences in subjective emotional experience between different categories of word pairs were taken into account in our subsequent analyses.

### Emotion-specific effects on encoding-related brain activity

To elucidate the neural mechanisms of memory modulation by emotion, we first identified brain regions involved in the successful encoding of word pairs related to different emotion categories: disgust (DIS), fear (FEA), neutral (NEU). Importantly, we controlled again for the level of arousal associated with each condition by including the mean difference in arousal ratings (see: Fig. 1e) for all word pairs as a covariate in the analysis. We focused our initial analyses on the activity within key anatomical regions of interest (ROIs) implicated in memory and emotion interactions, including the bilateral amygdala (AMY), perirhinal cortex (PRC), parahippocampal cortex (PHC), and hippocampus (HC). For each ROI and each emotion condition separately, we extracted activity during successful (later correctly recognized) encoding of word pairs. We found that these regions contributed differently to successful encoding [F(7,350) = 4.378, p = .001, η^2^ = .081] and that their engagement was dependent on emotion condition (interaction effect [F(14,700) = 5.522, p < .001, η^2^ = .099]). To unpack these effects, we run a post hoc analysis of simple effects restricted to the two emotion categories (DIS, FEA) which showed that successful encoding produced significantly higher activity for disgust than fear in left AMY (p < .001) and left PRC (p = .011). On the other hand, successful encoding activity was significantly higher for fear than disgust in right PHC (p = .007) and marginally in right HC (p = .059).

To complement the ROI analyses based on a priori hypotheses about their role in emotional memory formation, we also tested for any difference in large-scale cortical networks associated with successful encoding of different emotion conditions using a whole-brain random-effect analysis. We compared emotional to neutral conditions (EMO > NEU), and both emotion categories with one another (DIS > FEA and FEA > DIS).

Critically, mean ratings of arousal were again included as covariates to rule out any confound with emotion category effects (for a model before regressing out arousal, see Supplementary Fig. S3 and Supplementary Table S2). We found that in general, successful encoding of emotional pairs (EMO > NEU) engaged medial prefrontal and parietal regions. More specifically, however, successful encoding of disgust-related unitizations (DIS > FEA) was related to activations in left AMY, left INS, and left dmPFC, as well as other extensive clusters of frontal, temporal and posterior parieto-occipital regions, related for instance to precision of episodic memory (Richter, Cooper, Bays, & Simons, 2016a). Successful encoding of fear-related unitizations (FEA > DIS) showed activations in bilateral PHC, temporal and parietal regions (see Fig. 3 and Supplementary Table S3).

**Fig. 3.**
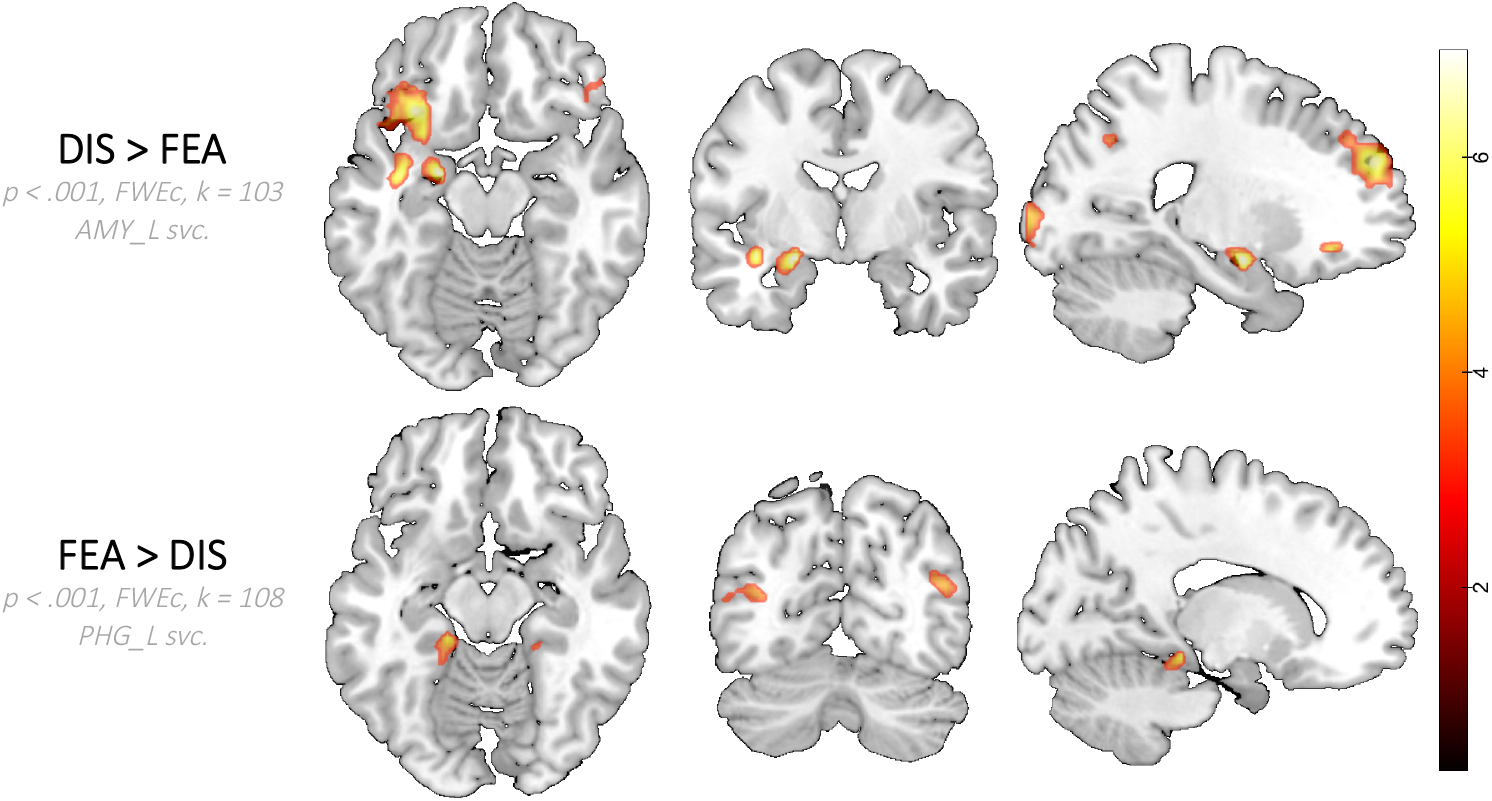
Differences in brain activation during successful encoding between basic emotion conditions (disgust and fear), after regressing out the effects of arousal; DIS – disgust, FEA – fear; colour scale represents a range of t-values; k – cluster extent, svc. – small volume correction, FWEc - voxel-wise height threshold of p < 0.001 (uncorrected) combined with a FWE-corrected cluster-level extent threshold of p < 0.05.

**Fig. 4.**
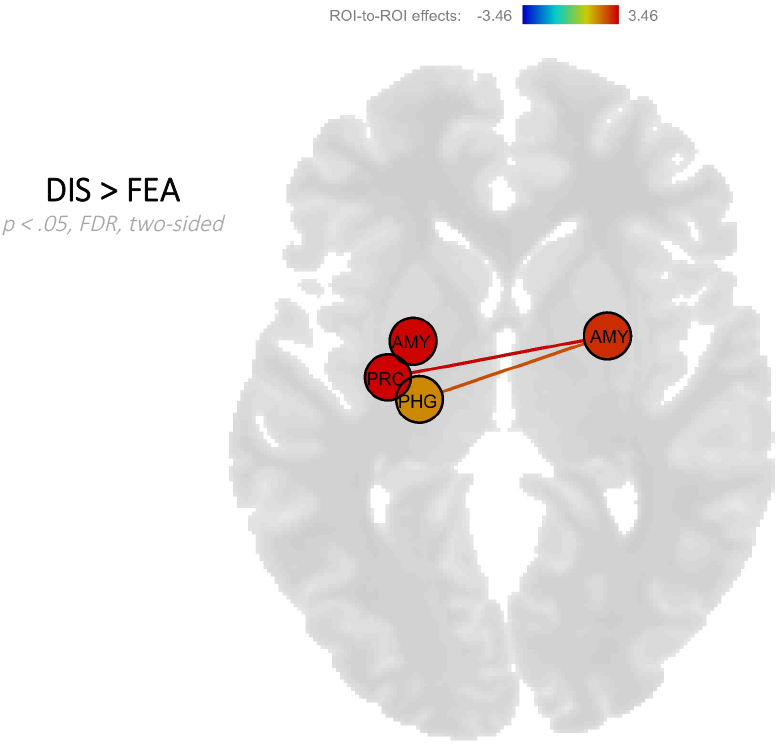
Increase in functional connectivity during successful encoding of old word pairs related to disgust when compared to fear; DIS – disgust, FEA – fear, pos. – positive, AMY – amygdala, PRC – perirhinal cortex, PHC – parahippocampal cortex; colour scale represents the range of effect sizes (regression coefficients); two-sided threshold of p < 0.05, FDR-corrected at the seed-level.

**Fig. 5.**
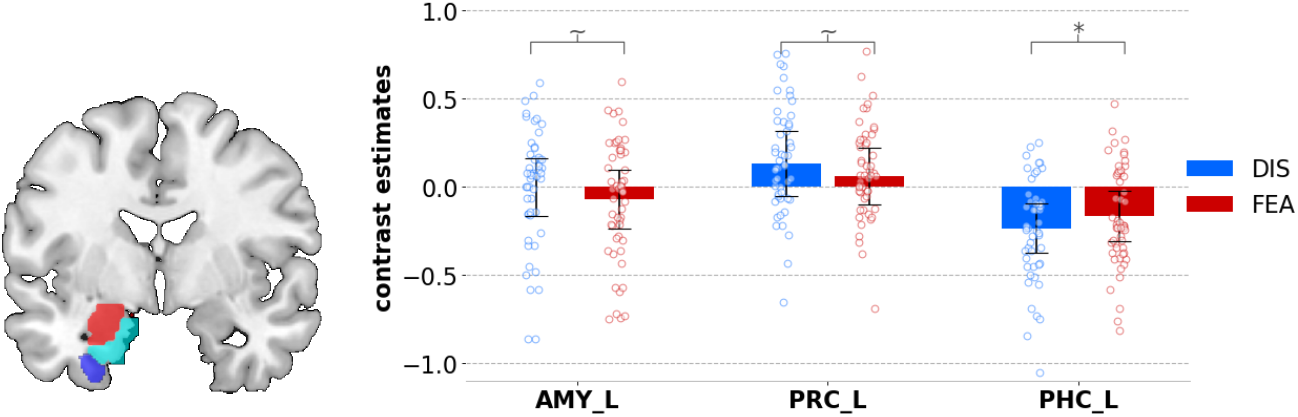
Emotion-related differences in recognition activity. Bars represent contrast estimates for correct recognition activity observed in presented anatomical ROIs (marked on the brain template: left AMY in blue, left PRC in cyan; left PHC in violet), split according to emotion factor (disgust- and fear-related word pairs); error bars represent one standard deviation, dots represent individual subjects’ scores; DIS – disgust, FEA – fear; ~ trend toward significance, * p < .05.

We also examined how these successful encoding effects were modulated by individual differences in experienced emotion using the affective ratings of word pairs collected in the third experimental session. Remarkably, brain activity during encoding was modulated by the disgust intensity in left dmPFC and bilateral AMY, whereas it was modulated by the fear intensity in bilateral dmPFC, and medial parietal regions, related for instance to threat contextualization (Bukalo & Holmes, 2018) and vividness of episodic memory (Richter et al., 2016a) (Supplementary Fig. S4 and Suppementary Table S4). No modulation of MTL regions was found for other affective parameters (impact, valence, and arousal).

Finally, the functional connectivity analysis was performed to determine any changes in temporal synchronization of the MTL regions, beyond the modulation of their activation level. To this aim, we used a gPPI analysis to analyze the interaction between functional connectivity of each source and target ROIs (ROI-to-ROI analysis) and experimental conditions (McLaren, Ries, Xu, & Johnson, 2012). Thus, our gPPI analysis treated these regions as seeds and determined how they were functionally coupled with the remaining ROIs in pre-defined contrasts of interest. Critically, when directly comparing disgust- and fear-related word pairs (DIS > FEA), we observed an increase in connectivity of the left AMY seed with left PRC [t(51) = 3.05; punc. = .004; pFDR = .026], and right AMY seed with left PRC [t(51) = 3.46; punc. = .001; pFDR = .008] as well as right AMY seed with left PHC [t(51) = 2.59; punc. = .012; pFDR = .043]. There were no results for the opposite contrast, meaning that no significant change in connectivity was found for the selected ROIs in the FEA > DIS contrast.

These results indicate that correct encoding of disgust-related word pairs was related to an increase in functional connectivity between brain regions typically related to affective processing, context memory, and items or unitized associations.

### Emotion-specific effects on recognition-related brain activity

Although it was not the main focus of our study, we also investigated brain activity during recognition and found that it was partly consistent with the findings from encoding phase. As previously, we controlled for the level of arousal associated with each condition by including the mean difference in arousal ratings (see: Fig. 1e) for all word pairs as a covariate in the analysis. We first focused our analyses on key anatomical regions of interest (ROIs) (bilateral AMY, PRC, PHC, and HC). We found that these regions contributed differently to correct recognition (main effect of ROI [F(7,350) = 6.266, p < .001, η^2^ = .111]) and that their engagement was dependent on emotion condition (interaction effect [F(14,700) = 2.301, p = .021, η^2^ = .044]). A post hoc analysis restricted to emotion categories (DIS, FEA) revealed that activity during correct recognition was marginally higher for disgust than fear in left AMY (p = .063) and left PRC (p = .061). On the other hand, activity was significantly higher for fear in left PHC (p = .024). No difference was observed in hippocampus.

Similar to encoding, we tested for concomitant differences in large-scale cortical networks using whole-brain random analysis to compare recognition activity between emotional conditions (EMO > NEU, as well as DIS > FEA and FEA > DIS). These comparisons were performed for old word pairs after excluding effects of the same contrasts for new word pairs, in order to isolate activity related to memory from perception of emotion-related words (Voss, Bridge, Cohen, & Walker, 2017). Again, we also regressed out arousal ratings in this analysis (Fig. 6), for a model without arousal regressed out, see Supplementary Fig. S5, S6 and Supplementary Table S7. A direct comparison testing for disgust effect on memory (old DIS > FEA masked exclusively by new DIS > FEA) revealed activations in frontal and parietal regions, typically associated with attentional control and semantic processing. For fear (old FEA > DIS masked exclusively by new FEA > DIS), we observed activations in parietal regions previously implicated in spatial orientation and visual imagery (Supplementary Table S8).

**Fig. 6.**
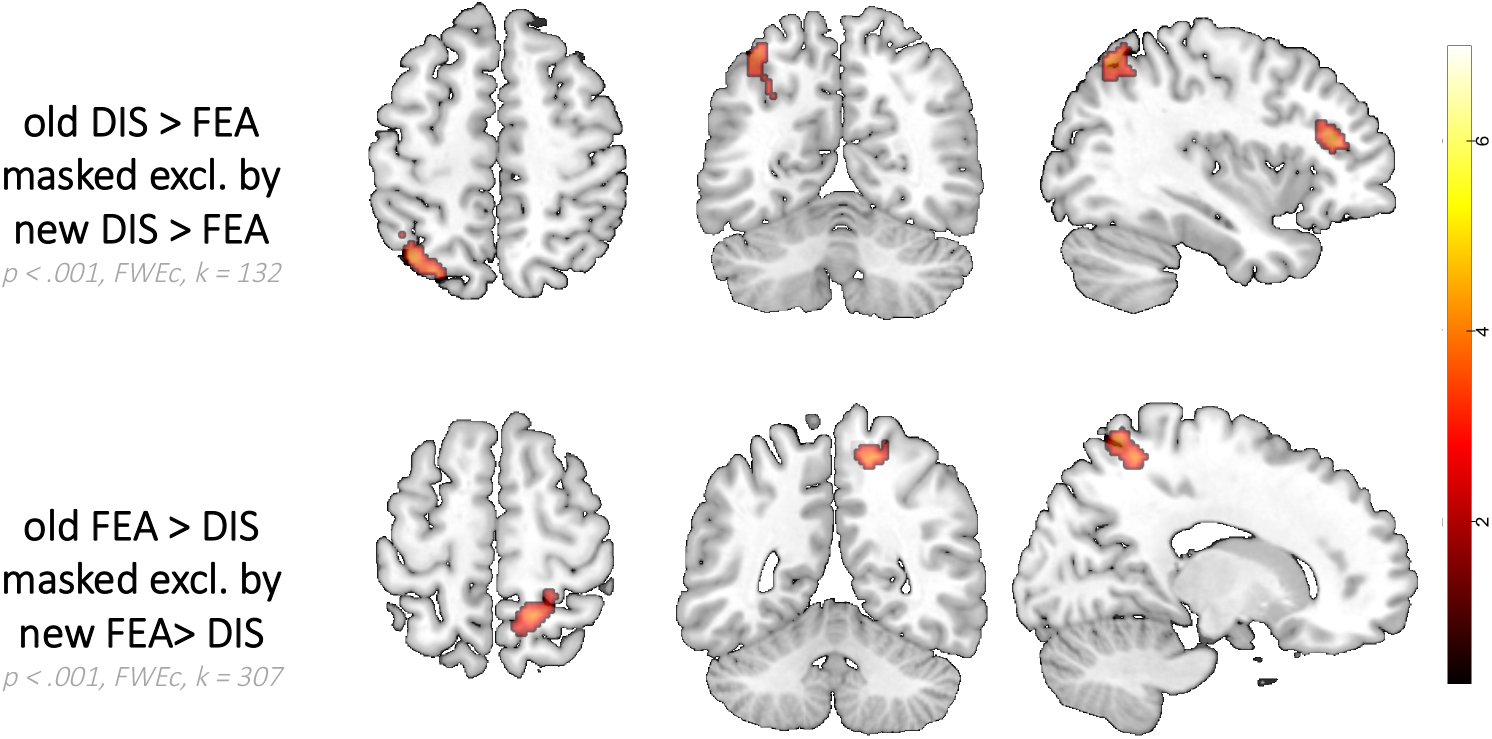
Differences in brain activation during successful recognition between basic emotion conditions (disgust and fear), after regressing out the effects of arousal. Effects for old word pairs were exclusively masked by the same effects for new word pairs; DIS – disgust, FEA – fear, corr – correct; colour scale represents a range of t-values; k – cluster extent; FWEc - voxel-wise height threshold of p < 0.001 (uncorrected) combined with a FWE-corrected cluster-level extent threshold of p < 0.05.

Additional results concerning the recognition activity as a function of individual ratings of experienced emotion are presented in supplementary Fig. S7 and Supplementary Table S9. The ROI-to-ROI functional connectivity analysis did not reveal any significant results.

### Encoding and reinstatement modulated by emotion

All analyses presented so far considered memory and emotion effects for encoding and recognition phases, separately. However, in line with current models of episodic memory, we aimed to investigate the role of emotion and AMY engagement also in trial-specific reinstatement of brain activity patterns.

To this aim, we performed a split-half correlation analysis within the framework of representational similarity analysis (RSA (Kriegeskorte, 2008)), using both the whole-brain searchlight and ROI methods (see: Methods). The searchlight analysis was performed with a sphere of interest moving from one voxel to the next through the whole brain volume (Kriegeskorte, Goebel, & Bandettini, 2006), whereas ROIs related to emotion and memory were defined anatomically, as in previous sections. Using this voxel-wise multivariate approach, we computed encoding-retrieval similarity (ERS) (Staresina, Henson, Kriegeskorte, & Alink, 2012) index, that is correlations between encoding and recognition patterns for matching versus non-matching trial pairs across experimental conditions. We found a significantly higher ERS index for all correctly than incorrectly (CORR > INCORR) recognized stimuli in left AMY and right PHC, and for disgust-than fear-related (DIS > FEA) correctly recognized word pairs in right PHC (both p < .01, k = 5, svc.).

However, a recent study (Xiao et al., 2017) demonstrated that the relationship between encoding and retrieval might be more complex than a faithful replay of past events and involve additional constructive, transformation processes. Therefore, complemented the previous analysis with reciprocal comparisons between encoding and recognition brain activity patterns reflecting more complex relationships, based on (Danker, Tompary, & Davachi, 2017). Specifically, we investigated how AMY engagement during encoding determines subsequent reinstatement. We first queried the *recognition* data for voxels in which a trial-by-trial univariate activity was predicted by (correlated with) trial-by-trial encoding activity in AMY. Second, we queried the *encoding* data for regions whose activation predicted further trial-by-trial reinstatement of activation patterns during recognition, as indicated by the ERS index in masks defined specifically for each emotion category.

In the first analysis, we found that activation of the left AMY during encoding significantly modulated neural activity observed during recognition in the right HC, left HC and right AMY (see Fig. 7 and Supplementary Table S10) regardless of emotion category.

**Fig. 7.**
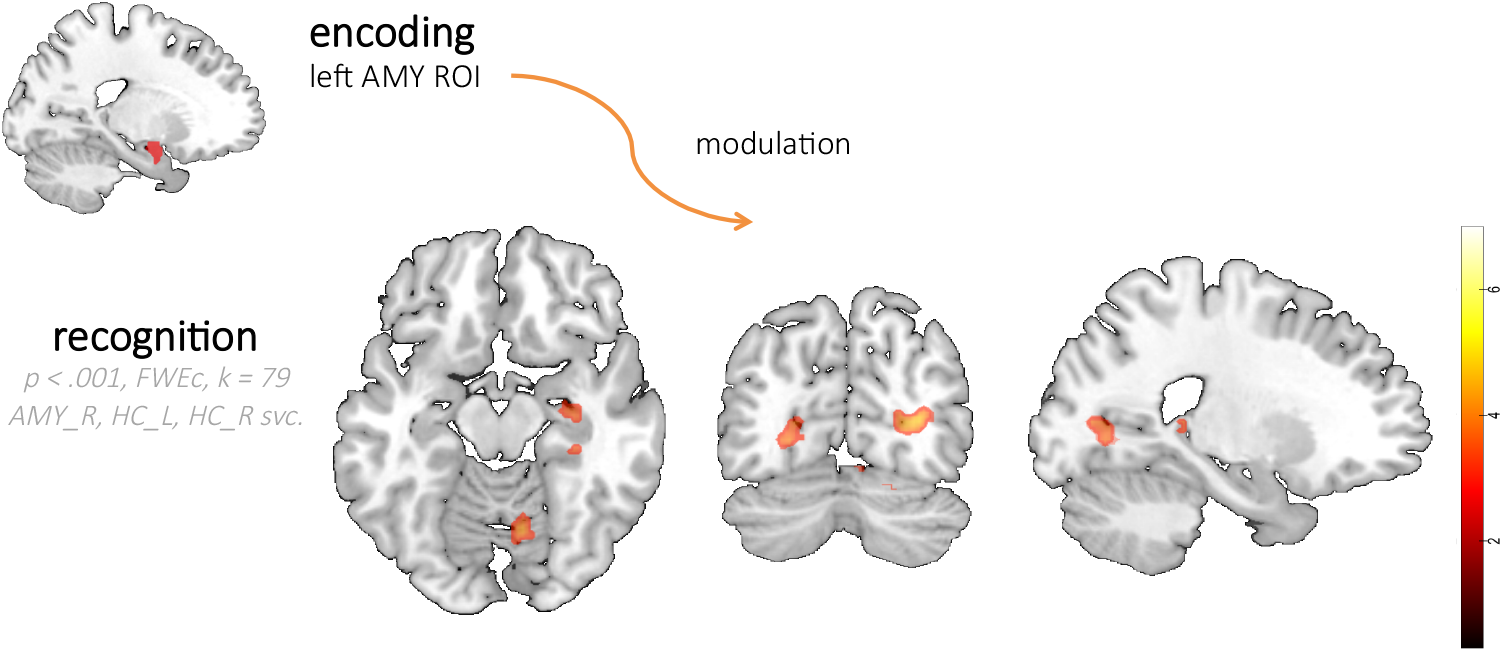
Brain activation during recognition, modulated by trial-by-trial activation in the left AMY ROI at encoding. AMY – amygdala, HC – hippocampus; FWEc – cluster-level FWE–corrected; k – cluster extent, svc. – small volume correction; colour scale represents a range of t-values.

In the second analysis, we found that during encoding of disgust as compared to fear (DIS > FEA), the activation in left AMY as well as other regions (Fig. 8 and Supplementary Table S6) predicted further ERS in emotion-specific ROIs.

**Fig. 8.**
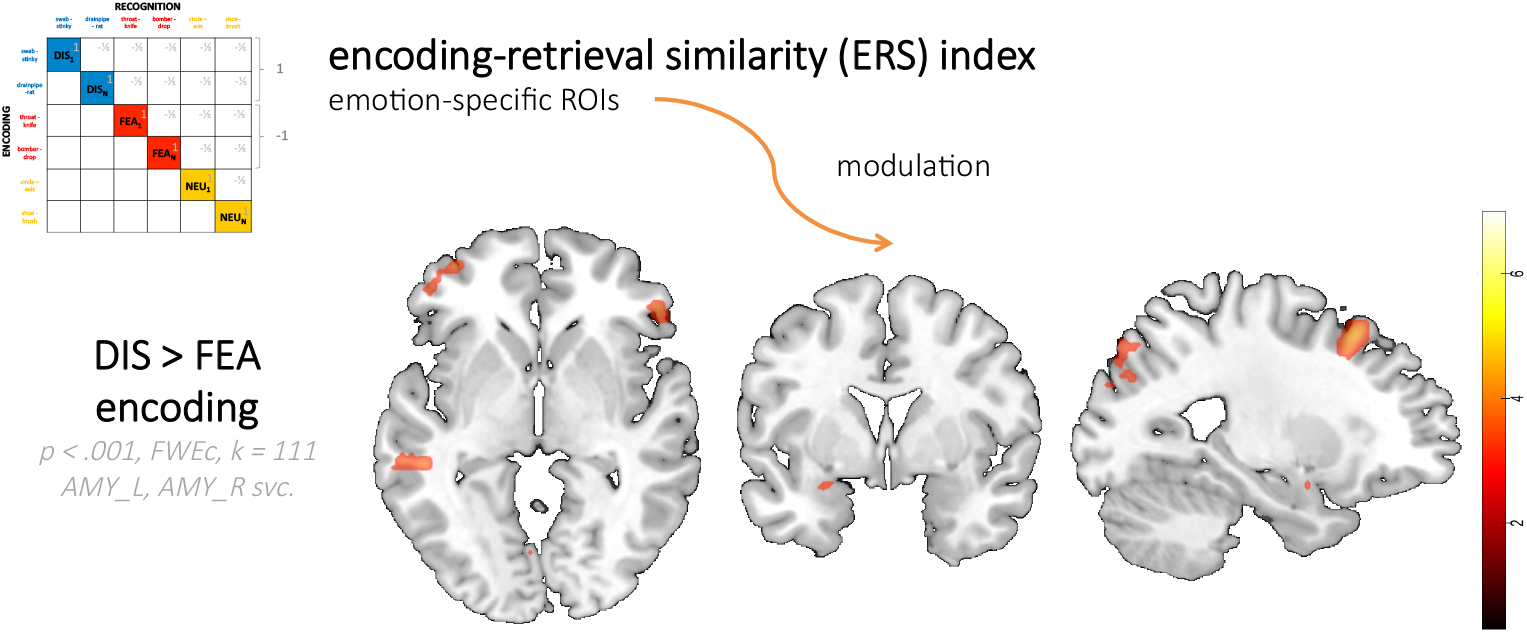
Brain activation during encoding modulated by trial-by-trial ERS. AMY – amygdala, IFG – inferior frontal gyrus; FWEc – cluster-level FWE–corrected; k – cluster extent, svc. – small volume correction; colour scale represents a range of t-values.

In total, these univariate and multivariate results show that AMY played an important role in the encoding and reinstatement of brain activity. Specifically, there was a positive correlation between the activity of left AMY during encoding and regions engaged in mnemonic and attentional processes during recognition. We also found a correlation between ERS index computed for each trial and left AMY activation on the corresponding encoding event, for disgust compared to fear. This suggests that the magnitude of initial AMY activation is related to the fidelity of subsequent reinstatement of activity patterns during recognition, particularly of disgust-associated stimuli. Consistent with current theories of episodic memory, these findings demonstrate a previously unknown critical link between AMY activation during encoding of an emotional experience and subsequent memory reinstatement.

## Discussion

How emotion influences episodic memory is a fundamental question for affective and cognitive neuroscience. While previous studies focused mainly on showing how mnemonic processes are modulated by the arousal evoked by to-be-remembered information, here for the first time we uncover brain mechanisms of differential encoding and reinstatement modulation by discrete emotion categories, independent of valence and arousal. Instead of separating emotion effects on item and associative memory, we investigated a specific type of very close associations under unitization instructions that might be processed as single items. Critically, we complement previous studies findings on emotion effects selectively during encoding or retrieval by studying how emotion influences memory reinstatement.

Our results revealed that after 2-3 weeks, word-pairs evoking disgust were remembered better than word pairs evoking fear, an effect reflected by different brain pathways. Specifically, univariate activation in AMY and PRC supported better memory for disgust-related word pairs, whereas univariate activation in PHC and HC supported better memory of fear-related word pairs. We also found that AMY was temporally synchronized with other MTL regions during successful encoding of disgust more than fear. Critically, AMY activity at encoding was shown to modulate brain activation in HC at recognition, and to predict a subsequent fidelity of memory reinstatement of emotion-specific representations of individual word pairs, more for disgust than fear. These results unveil how different emotion categories such as disgust and fear differentially modulate memory despite similar negative valence and arousal level. Finally, our findings uncover a previously unknown, emotion-specific role of amygdala for successful encoding and reinstatement.

### Differences in memory modulation by disgust and fear

Our first major finding was that disgust and fear-related information produced behaviourally distinct effects on long-term memory of unitizations. Even though these two emotions are negative, arousing, and associated with motivational avoidance (Krusemark & Li, 2011), they are known to induce distinct physiological responses (Critchley et al., 2005) and different effects on cognitive processes including item memory. Better recall of disgust-compared to fear-related stimuli was previously demonstrated for single words (Charash & McKay, 2002) and images (Croucher et al., 2011). Here we show better memory for unitized word pairs evoking disgust compared to fear, an effect unexplained by differences in arousal level. Based on previous literature, we assume that this effect is unlikely to reflect a differential influence of disgust and fear on attentional processes (Chapman, 2018; Chapman et al., 2012; van Hooff et al., 2014). Also, we did not find any differences in cognitive effort (measured as RT) during recognition among these two basic emotion categories. Given that we investigated the influence of disgust and fear on memory of verbal associations, we additionally controlled for other possible confounds: imageability success during encoding, and semantic coherence of word pairs (Supplementary Fig. S2). No differences between disgust- and fear-related word pairs were found for these factors. A recent study attributed the higher level of free recall for disgust-related than neutral and fear-related words to deeper elaborative encoding (Ferré, Haro, & Hinojosa, 2018). Since the nature of our task (creating common mental representations of word pairs) forced elaborative processing for all the experimental conditions, it should eliminate also such differences. Altogether, disgust appears to have a special salience in memory for verbal associations regardless of possible confounding factors.

Extending our behavioural effects, the analyzed neuroimaging data revealed that disgust and fear engaged distinct neural pathways of mnemonic processes. First and foremost, we observed a specific involvement of AMY in encoding and reinstatement of disgust-related verbal unitizations. In animals, AMY activity was shown to initiate autonomic responses to salient stimuli (LeDoux, Iwata, Cicchetti, & Reis, 1988), consistent with later human neuroimaging studies (Critchley, Mathias, & Dolan, 2002). Despite a traditional focus on the processing of fear and threat in AMY (Phelps & LeDoux, 2005), recent studies demonstrate that AMY may activate to any stimuli characterized by a high level of personal impact, independent of intrinsic emotional properties (Ewbank, Barnard, Croucher, Ramponi, & Calder, 2009), relevance detection (Sander, Grafman, & Zalla, 2003) and evaluative processing of goals (Cunningham, Van Bavel, & Johnsen, 2008). Here we found increased AMY activation for the memory of word pairs related to disgust, which were indeed characterized by higher subjective ratings of impact, but not arousal. Also, we show that individual ratings of disgust, but not ratings of arousal or impact, modulated encoding activity in AMY.

Our results are in line with previous studies showing that AMY activity at encoding increases the likelihood of remembering emotional, but not neutral items (Dolcos, LaBar, & Cabeza, 2005), even without a conscious emotional experience (Inman et al., 2018). However, the current study is the first to show that AMY differentially contributes to memory for two distinct negative emotions, even after controlling for arousal level (de Voogd, Fernandez, & Hermans, 2016). Most importantly, we also found that left AMY activation during encoding critically influenced the fidelity of brain activity pattern reinstatement during recognition, as shown by our univariate and multivariate analyses. An important point of reference comes from animal studies, where AMY coordinated with HC (Girardeau, Inema, & Buzsáki, 2017), or even AMY itself (Reitich-Stolero & Paz, 2019) was shown to play a crucial role in reactivation or rehearsal of contextual affective memories, as a mechanism of enhanced consolidation. Previous human studies employing multi-voxel analyses have only investigated the relationship between encoding and retrieval brain activation pattern (Xue, 2018), either for emotional items (Ritchey, Wing, Labar, & Cabeza, 2013) or neutral associations (Danker, Tompary, & Davachi, 2016), but to our knowledge emotion modulation of episodic memory has never been tested this way.

Another brain structure playing a crucial role in memory of disgust was PRC. Other findings support complex multi-modal item representations in this region (Ranganath & Ritchey, 2012), such as integration of novel odors with visual category information (Qu, Kahnt, Cole, & Gottfried, 2016), or integration of visual with conceptual object features (Martin, Douglas, Newsome, Man, & Barense, 2018). PRC was also shown to reflect semantic similarity between words (Bruffaerts et al., 2013), yet here we found no difference in the semantic coherence of word pairs between disgust and fear. This region has direct connections and a privileged access to signals from AMY (Pitkänen et al., 2000), which may be important for coding item salience. Greater PRC activity during emotional compared to neutral memory encoding was observed in previous research (Dolcos, LaBar, & Cabeza, 2004), but only if it was accompanied by better recollection compared to neutral items, similar to our study. Also, the engagement of PRC might result from the multisensory nature of disgust processing, although this factor was not manipulated nor quantified in our experiment.

### Differences possibly related to unitization process

We found that bilateral PHC and right HC played a crucial role in successful encoding of word pairs evoking fear, again regardless of arousal levels. The ROI analyses showed that the right PHC and right HC were activated more during encoding of fear-than disgust-related word pairs, whereas left PHC activated during recognition of fear-more than disgust-related word pairs. Given that PHC is typically related to context processing (Murray & Kensinger, 2013; Yonelinas & Ritchey, 2015), these results may suggest that fear-related word pairs might evoke pre-existing associations related to the meaning of words within pairs, but be less efficiently associated together by unitization. However, our measure of integrative imageability success during encoding did not reveal any difference between emotions. In any case, the current results clearly converge to indicate that different unitization and memory mechanisms were engaged when modulated by disgust- and fear-related word pairs.

Previous studies have indicated a double dissociation between PRC and PHC for encoding of object vs. scene context (Staresina, Duncan, & Davachi, 2011), paralleling the dissociation found here between disgust and fear. One possibility is that fear-related unitized mental representations were processed in a scene-like fashion of spatial exploration, in line with an effect of threat signals on the monitoring of the environment. Another possibility is that emotions may generally enhance context processing, particularly in the case of fear. PHC could therefore mediate this strong connection between contextual processing and emotional information, facilitating emotion understanding but also subsequent episodic memory (Aminoff, Kveraga, & Bar, 2013).

The involvement of HC in emotional memory has been also intensely investigated. One important insight to this issue was provided by a study (Dandolo & Schwabe, 2018) where superior memory for emotional material came at the cost of reduced memory for contextual details. This was reflected by an increase in functional connectivity between the left HC and right AMY for negative compared to neutral photographs. Another study(Madan et al., 2017) reported that associative memory for negative information was impaired, but accompanied by increased activity in AMY, as well as in the HC. The authors suggested that when sufficiently arousing information prevent unitization-based encoding supported by other MTL-cortex regions, an alternative, relational HC-dependent encoding strategy may be engaged. Our results showed that AMY activity during encoding modulated the activity of right HC during recognition, shedding a new light on this relationship also across the two key stages of mnemonic processes, i.e., encoding and recognition.

It is important to note that our results conflict with previous reports of a disruptive role of AMY during encoding of arousing associations (Bisby, Horner, Horlyck, & Burgess, 2016; Madan et al., 2017; Murray & Kensinger, 2014a; Okada et al., 2011). However, they align with recent finding that the engagement of both AMY and PRC can support the recollection advantage for emotional items (Ritchey et al., 2019), which reflect how unitized word pairs were memorized and recognized. Our results showed an increase in functional connectivity between AMY and PRC for disgust compared to fear. Despite no differences in the strength of associations created during encoding at the behavioural level, the interplay between AMY and PRC may be promoted by a more successful unitization (Madan et al., 2017). Also, our data converge with the *emotional binding* account (Yonelinas & Ritchey, 2015), according to which AMY mediates the recollection of item-emotion bindings that are forgotten more slowly than item-context bindings supported by the HC (Yonelinas & Ritchey, 2015). This account is in line with our results whereby more AMY activation related to encoding and recognition of disgust (as compared to fear-related and neutral word pairs) induced better memory. A question remains, however, if item-emotion associations can be unitized, as instructed in our experiment. If they can, the unitized item-emotion associations should be supported by AMY and PRC, as well as later recognized based solely on familiarity. However, we tested only recognition memory and not associative memory, so this prediction will require further studies.

### Possible explanations of distinct brain mechanisms

Although disgust and fear modulated memory of unitized associations through partly distinct neural mechanisms, we do not imply that there exist separate neural substrates for basic emotion categories, which has long been debated (Saarimäki et al., 2018). Rather, our study was designed to determine whether, and how, encoding and reinstatement of verbal associations may be differently modulated by disgust and fear regardless of overall negative valence and high arousal. Above all, we do not imply that AMY is not involved in fear processing (Hermans et al., 2017; Vuilleumier, Armony, Driver, & Dolan, 2001) or that PHC cannot be involved in associative memory of other emotion categories (Aminoff et al., 2013). However, we surmise that fear and disgust might recruit partly distinct emotion appraisal components that activate different kind of associations in memory and thus recruit partly distinct neural systems holding such associations.

We should however acknowledge some limitations to these conclusions. It is possible that this dissociation is specific to verbal material, which is more likely to evoke disgust than fear. Thus, a different pattern might be observed for other modalities, for instance with visual or auditory material (Artur Marchewka et al., 2016; Zimmer, Höfler, Koschutnig, & Ischebeck, 2016). AMY activation can influence both visual and semantic processing (Skipper, Ross, & Olson, 2011), but how these influences differ for associations has not been explored.

It is also possible that our effects were specific for this kind of memory task, with unitization instruction during encoding and recognition after a long-term delay, which may require complex cognitive operations (Memel & Ryan, 2018; Tibon, Greve, & Henson, 2018). Moreover, our experimental design does not allow us to draw any conclusions about emotional modulation of associative memory (Ritchey, Wang, Yonelinas, & Ranganath, 2018), so we cannot determine whether our participants recollected information based on recollection or familiarity and how continuous is the unitization process across these conditions (Ritchey, Montchal, Yonelinas, & Ranganath, 2015).

Also, our experimental design was limited to comparing only disgust, fear and neutral content, preventing us to generalize our findings to other basic emotion categories, either negative or positive (Diano et al., 2017). Last but not least, it must be underscored that studies on memory and emotion exhibit a wide variety of experimental procedures, with most manipulating only valence and arousal, such that it is difficult to directly compare them (Dolcos et al., 2017).

### Conclusions

In sum, our results reveal remarkable differences in memory modulation between disgust and fear, both at the behavioural and neuronal level. We found not only that long-term recognition memory for unitized word pairs is modulated by emotion, but this enhancement appears stronger for disgust-than fear-evoking material, an effect unexplained by arousal level. Greater AMY and PRC activations accompanied successful memory of disgust-related word pairs, whereas PHC and HC supported better memory for fear. Moreover, the critical AMY activation during encoding was correlated with univariate activation in HC during retrieval, as well as with memory reinstatement of trial-by-trial emotion-specific representations. This correlation was higher for disgust than fear, emphasizing the dissociation between these two essential emotions. These differences might result from distinct associative processes at play, or particular motivational and physiological responses evoked by disgust and fear. For instance, fear-motivated avoidance may protect one from direct danger, disgust-motivated avoidance might more often be linked to anticipatory sensation or imagery (see Woody, 2000).

To conclude, we demonstrate that there might be distinct neural pathways engaged in memory modulation for different emotion categories, over and above valence and arousal dimensions. Finally, we shed a new light on the role played by amygdala during encoding and reinstatement of memories related to specific emotions. These results provide important insights for emotion and memory research, as well as potential interventions.

## Supporting information

Supplementary tables encoding

Supplementary tables recognition

## Materials and methods

### Participants

Fifty-nine native Polish speakers (29 F and 30 M; aged 20-33; M = 24,81, SD = 3.14) without history of any neurological disorders or treatment with psychoactive drugs, right-handed, with normal or corrected-to-normal vision, gave written informed consent and participated in the study. They were mostly students and professionals living in Warsaw, with at least secondary education. Six subjects were excluded from the experimental group due to technical problems during experimental procedure and one additional subject did not take part in the second experimental session. Thus, behavioural and neuroimaging data collected from fifty-two subjects was further analyzed (24 F and 28 M; aged 20-33; M = 24,83, SD = 3.21). The local Research Ethics Committee at Faculty of Psychology, University of Warsaw approved the experimental protocol of the study.

### Stimuli

360 words were selected from the Nencki Affective Word List (NAWL) (Riegel et al., 2015). According to the collected affective ratings and a novel method of classification based on Euclidean distances (Małgorzata Wierzba et al., 2015), a total of 120 words eliciting disgust, 120 words eliciting fear, and 120 neutral words was selected. As presented in Fig. 1, the stimuli were counterbalanced on all the other affective scales (valence, arousal, and intensities of other basic emotions). Subsequently, in each emotion category, 60 word pairs (6-25 characters) were formed in order to limit possible associations and suggest a specific semantic meaning. The word pairs consisted of emotionally and semantically congruent words (M = 3.61, SD = .65) as rated by 5 independent raters (6-point scale, with −1 = words are semantically opposite, 0 = words are not semantically congruent and 5 = words are highly semantically congruent). Thirty-five out of 60 word pairs in each emotion category (disgust, fear, neutral) were used in both encoding and recognition sessions as targets, whereas 25 were additionally included as lures to the recognition session. The final set of word pairs can be illustrated with the following examples (translated to English, also in the figures) for disgust: *swab - stinky, drainpipe -rat, shit – pigeon*; for fear: *throat - knife, bomber - drop, truck – collision*; and for neutral state: *circle - axis, word - Latin, facade - stone*. The full list of Polish word pairs in the order of presentation during encoding and recognition sessions is included in the Supplementary materials, Table S1.

**Fig. 1.**
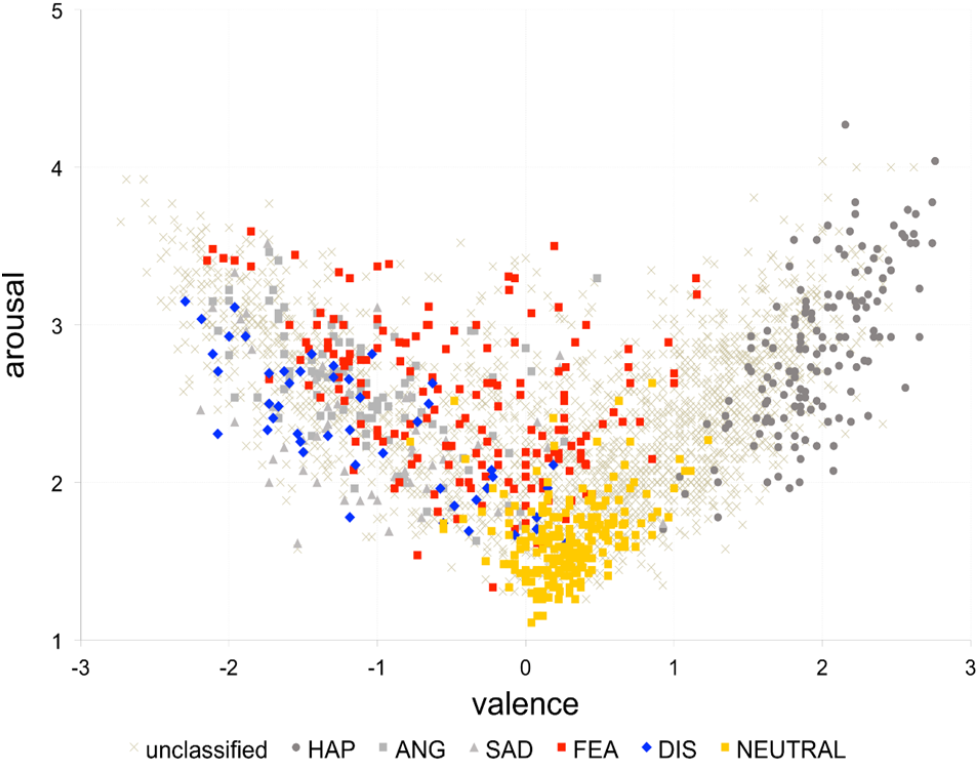
Distribution of the NAWL words assigned to experimental conditions of basic emotions (disgust, fear and neutral) in the affective space of valence and arousal. Each dot represent a single word, blue – disgust, red – fear, yellow – neutral; HAP – happiness, ANG – anger, SAD – sadness, FEA – fear, DIS - disgust (dataset available as an interactive browser: http://exp.lobi.nencki.gov.pl/nawl-analysis)

### Study design and experimental paradigm

The study consisted of three experimental sessions, two of which (encoding and recognition) were conducted with the use of MRI scanner, with a delay period of 15-19 days (M = 16.86, SD = 1.22) (Fig. 2). After the second session and a short break, the third experimental session started during which subjective affective ratings were collected through a web application running on a local server (http://exp.lobi.nencki.gov.pl/). All the experimental sessions took place in the Laboratory of Brain Imaging, Nencki Institute of Experimental Biology in Warsaw, Poland.

**Fig. 2.**
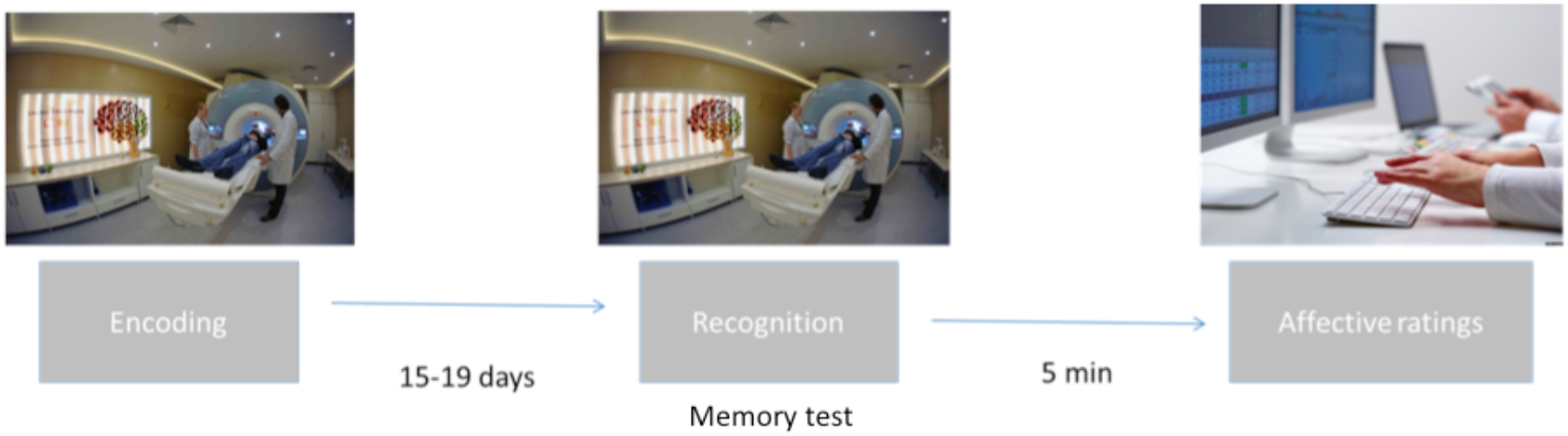
Overview of experimental procedure.

Before entering the scanner, subjects were given the details of scanning procedure and performed a brief practice session of memory task (comprising 10 word pairs) in a mock MRI scanner. The experimental procedure was programmed using the Presentation software (Neurobehavioral Systems, Inc., Albany, CA, USA) and displayed on an MR-compatible high-resolution LCD monitor positioned at the back of the scanner. Subjects observed the stimuli through the mirror system placed on the MR coil. The order of experimental trials in both sessions was pseudorandom under the following constraints: no more than 3 consecutive trials of the same emotion category (disgust, fear, neutral), no more than 3 of the same part of speech (noun, verb, adjective), no more than 4 old (targets) or new (lures) in the recognition session, and a maximized difference in semantic congruency (as rated by independent raters) of each two consecutive word pairs. In order to avoid the serial-position and recency effects, word pairs selected for both encoding and recognition sessions were divided into three parts (A, B, C), and presented in three variants of their order, counterbalanced across all the participants (encoding: ABC n = 21, BCA n = 20, CAB n = 18; recognition: ABC n = 20, BCA n = 20, CAB n = 19).

The first experimental session was encoding, as depicted in Fig. 3. The participants were presented (4s + jittered fixation cross for 3-7s) with 105 word pairs (35 disgust-related, 35 fear-related and 35 neutral), instructed to imagine a single mental representation for each word pair and memorize as many of them as possible. The participants were presented with all the word pairs twice in order to strengthen the memory trace. The second presentation took place during a separate scanning session, right after the first one, in an identical order. During the second presentation, each word pair (3 s + jittered fixation cross of 3-7s) was followed by a rating of integrative imageability success (Murray & Kensinger, 2014b), i.e. how successful a participant was when trying to imagine a single mental representation for a particular word pair (3s; 4-point scale, with 1 = not successful and 4 = very successful). The instruction was formulated as follows: “Try to memorize as many pairs of words as possible. In order to do so, imagine each two words as one, in a single mental representation. You will be presented with the pairs twice, in the same order. After the second presentation, you will be asked to rate the extent to which you can integrate each word pair into a single mental image (1-4 scale).” If not stated differently, fMRI data from both presentations are included in a single model in the presented analyses.

**Fig. 3.**
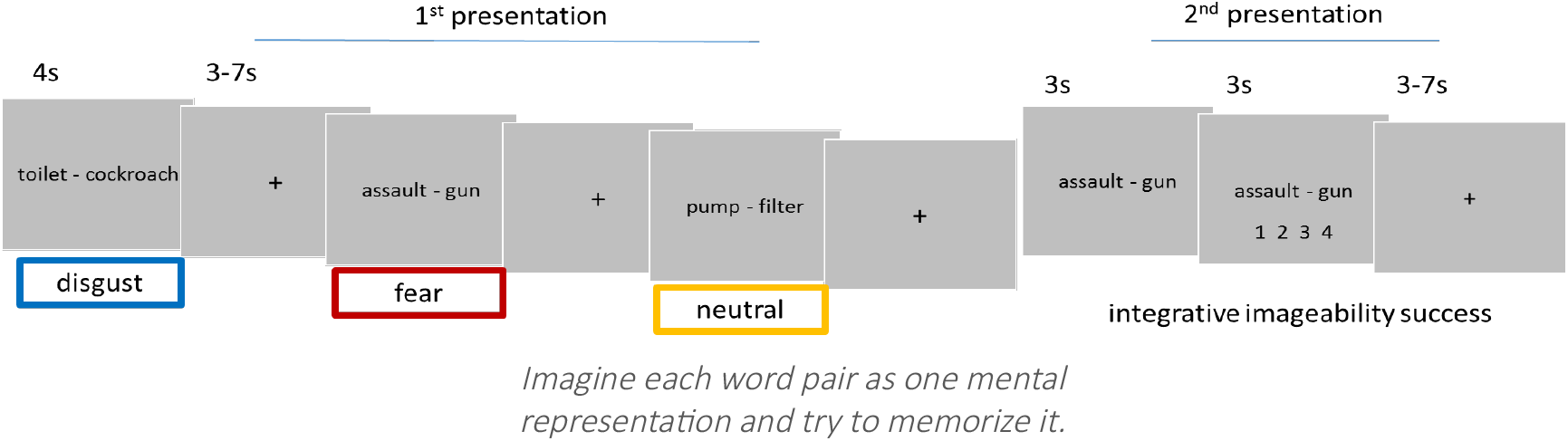
Example trial structure for all the emotion conditions during encoding.

After 15-19 days, the subjects took part in the recognition session (Fig. 4). During an item recognition task, participants were presented with 105 word pairs from the encoding list and 75 other word pairs as lures, that is 180 word pairs in total (2s + jittered fixation cross for 3-7s), and asked to determine (3s) whether a word pair was old (studied earlier) or new (not studied before). Additionally, participants were asked to indicate if they were sure or unsure of their responses (2s). The instruction was formulated as follows: “You will be presented with word pairs expressing different emotions or neutral. Your task will be to indicate if you can remember having seen them [O – old] or you cannot remember it [new – N], and if you are sure and remember your specific mental representation (Slotnick & Schacter, 2004) [S – sure] or you are unsure and do not remember your specific mental representation [U – unsure]. The recognition session was divided into two runs with a short break in between, for the comfort of participants. The fMRI data from both runs are included in a single model in the presented analyses.

**Fig. 4.**
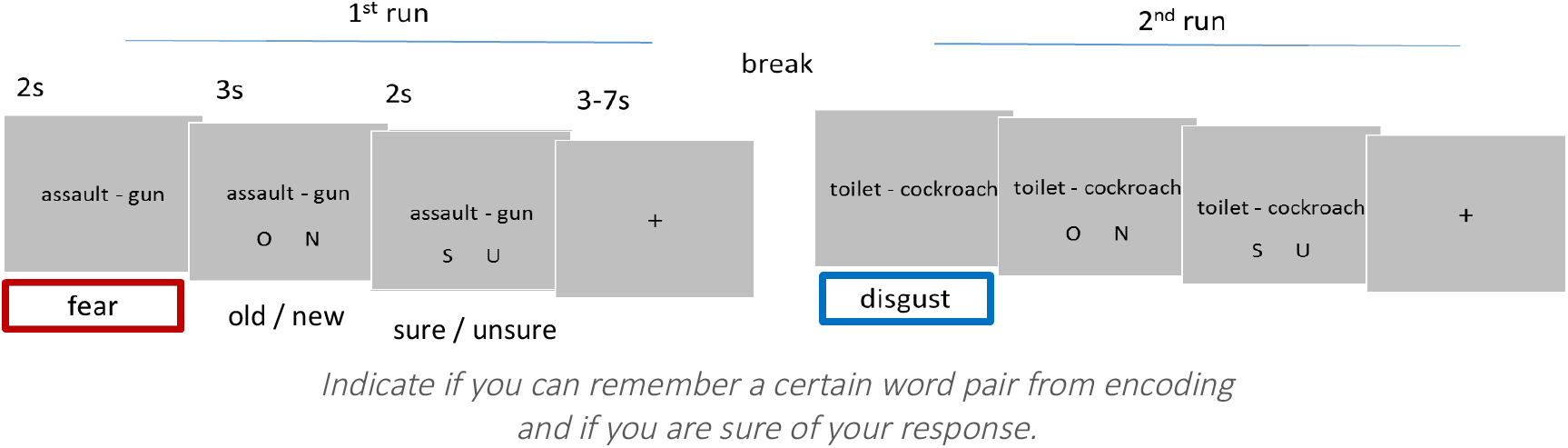
Example trial structure for all the emotion conditions during recognition.

In order to further control the individual differences in affective processing, after fMRI scanning the subjects took part in the third experimental session of affective assessment (Fig. 5). During this session, each participant was presented with the instructions, and subsequently with all the 180 word pairs used during encoding and recognition sessions. The task was to assess emotional properties of each word pair on several affective scales. Each word pair was presented in a full-screen view for 1 s. Then, the first scale of impact (Ewbank et al., 2009) appeared. The participants were asked to consider each word pair as a whole and assess whether they felt that its meaning created an instant and personal sense of impact on them (1 for very low and 9 for very high). Following, the assessed word pair appeared in smaller font in the upper part of a new screen, next to the remaining rating scales: an intensity of evoked disgust and fear (1 for not at all and 7 for very much), valence (−3 for negative, 0 for neutral and 3 for positive), and arousal (1 for unaroused and 5 for aroused, additionally accompanied by the self-assessment manikin, SAM, Lang et al., 1980). The screen remained visible to the participants until they completed all the ratings and pressed the “Next” button. Completing the whole affective assessment session took approximately 30 min.

**Fig. 5.**
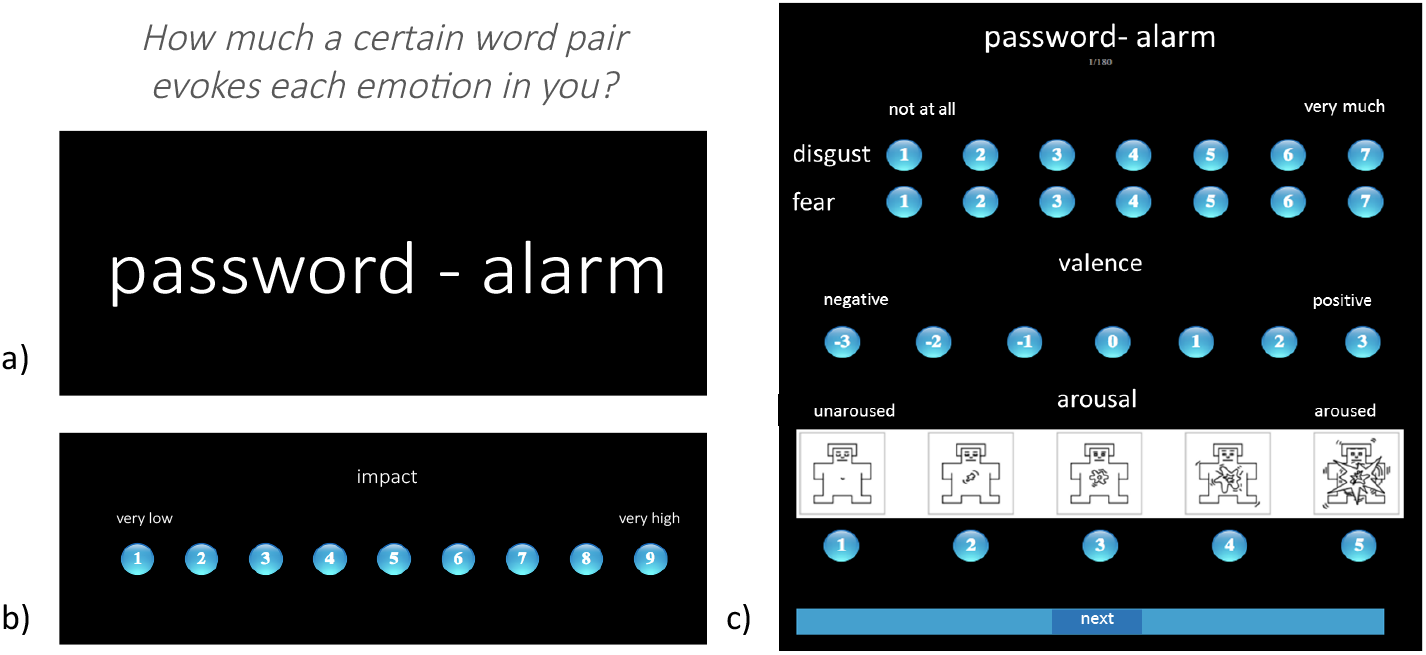
Example trial structure with a) first presentation of a word pair, b) scale of impact, c) all the other affective scales.

### Behavioral data analysis

First, memory for emotional and neutral stimuli was assessed as the proportion of correct responses in the recognition test. To analyze memory as a function of emotion category (3 levels: disgust, fear, neutral) and novelty (2 levels: old, new) as within-subject factors, repeated-measures (rm) analysis of variance (ANOVA) was performed on the recognition rate (proportion of correct responses) data. All the reported ANOVAs were performed with the Greenhouse-Geisser adjustments to the degrees of freedom when the assumption of sphericity was violated. Post hoc comparisons were performed using the Bonferroni adjustment for multiple comparisons.

Since the subjects had to make memory decisions under conditions of uncertainty and could potentially use certain strategies of responding, signal detection theory (SDT) (Wickens, 2002) was also applied to the collected data. Apart from the proportion of correct responses, that is *old* recognized as *old* (hits), the proportion of *new* incorrectly recognized as *old* (false alarms), the proportion of *new* correctly recognized as *new* (correct rejections) and the proportion of *old* incorrectly recognized as *new* (misses) were calculated. Based on these proportions, a sensitivity index (d’) was calculated to provide a measure of the separation between the means of the signal and noise distributions, compared against the standard deviation of this distribution: Z (hit rate) – Z (false alarm rate). It was calculated separately for each emotion category (disgust, fear, neutral). Finally, rm ANOVAs were performed on the obtained d’ values to test the effect of emotion (3 levels: disgust, fear, neutral) as a within-subject factor, on memory performance.

Next, in order to eliminate the possible confounds and assess the effect of emotion on memory performance more accurately, additional factor was included in the analysis. It has been long debated whether theoretical approaches of affective dimensions (Osgood & Succi, 1957) and basic emotions (Ekman, 1992) are mutually exclusive or complementary (Briesemeister, Kuchinke, & Jacobs, 2014) in explaining emotional experience. Therefore, we aimed at explaining whether any potential effect of basic emotions on memory could be explained by an associated level of arousal. Repeated-measures analysis of covariance (ANCOVA) was performed on d’ with emotion (2 levels: disgust, fear) as within-subject factor, with a covariate of the difference in mean level of arousal between disgust-related and fear-related stimuli (for the details of this difference, see further analyses of affective ratings).

Reaction times (RT) in the memory task performance were also analyzed to control for any differences in task difficulty (Theios, 1973) due to emotion. Recently, it has been also indicated that reaction times provide similar information to explicit ratings of memory confidence in their ability to quantify recognition memory decisions (Weidemann & Kahana, 2016). We performed rm ANOVA with emotion (3 levels: disgust, fear, neutral), novelty (2 levels: old, new) and correctness (2 levels: correct, incorrect) as within-subject factors.

The next research question was whether emotion effects are related to the strength of integration of encoded pieces of information or to the subjective strength of memory trace during recognition (Murray & Kensinger, 2014a; Richter et al., 2016a). To answer these questions, we analyzed additional responses given during encoding and recognition sessions concerning the integrative imageability success and the sureness of recognition responses, respectively. First, we performed rm ANOVA concerning the level of integrative imageability success, with emotion (3 levels: disgust, fear, neutral) and correctness (2 levels: correct, incorrect) as within-subject factors (only old word pairs were analyzed, as the integrative imageability success was rated during encoding session). Second, we performed rm ANOVA of the percentage of sure responses among correct responses, with emotion (3 levels: disgust, fear, neutral), and novelty (2 levels: old, new) as within-subject factors.

Individual affective ratings were used to analyze possible differences between word pairs assigned to respective emotion categories (disgust, fear, neutral), as an experimental manipulation check. Mean impact, disgust, fear, valence and arousal ratings were calculated within at the subject-level, and rm ANOVAs were performed with emotion (3 levels: disgust, fear, neutral) as within-subject factor for all the affective parameters. Paired t-tests for dependent samples were also performed to directly compare the mean affective ratings between disgust- and fear-related stimuli.

Finally, experimental manipulation was checked in terms of semantic congruency between words forming word pairs selected for the experiment, as rated initially by the independent raters. The mean semantic congruency for each word pair was compared between emotion categories (3 levels: disgust, fear, neutral) with one-way ANOVA to indicate if it was equal for all the experimental conditions.

### fMRI data acquisition and preprocessing

Magnetic resonance data was acquired on a 3T MAGNETOM Trio TIM system (Siemens Medical Solutions) equipped with a whole-head 32-channel coil. The following images were acquired within a single subject scanning during encoding session: structural localizer image, structural T1-weighted (T1w) image (TR: 2530 ms, TE: 3.32 ms, flip angle: 7°, PAT factor = 2, voxel size 1 × 1 × 1 mm, field of view 256 mm, volumes: 1), field map (TR: 488 ms, TE1: 5 ms, TE2: 7.46 ms, flip angle: 60°, voxel size 3 × 3 × 3 mm, field of view 216 mm, volumes: 1), first series of functional EPI images (TR: 2500 ms, TE: 30 ms, flip angle: 90°, PAT factor = 2, voxel size 3 × 3 × 2.5 mm, field of view 216 mm, volumes: 386 in encoding run 1, and 470 in encoding run), and second series of functional EPI images (same parameters, volumes: 2441 in recognition run 1, 441 in recognition run 2). The whole scanning session during encoding took approximately 37 minutes, and during recognition – approximately 40 minutes, independent of participants’ speed of responding as trials had a fixed duration.

At the initial step of fMRI data preprocessing, the DICOM series were converted to NIfTI using the MRIConvert (ver. 2.0.7; Lewis Center for Neuroimaging, University of Oregon). Then, the brain imaging data was preprocessed using Statistical Parametric Mapping (SPM12; Wellcome Department of Cognitive Neuroscience, University College London, London, UK) running under Matlab 2013b (Mathworks, Inc., Natnick, MA, USA). The preprocessing of data started with the correction of functional images for distortions related to magnetic field inhomogeneity and correction for motion by realignment to the first acquired image. Then, structural (T1w) images from single subjects were co-registered to the mean functional image and segmented into separated tissue classes (grey matter, white matter, cerebrospinal fluid) using the default tissue probability maps (TPM). Structural and functional images were normalized to the MNI space and resliced to preserve the original resolution. Finally, functional images were smoothed with the 6 mm FWHM Gaussian kernel. Unsmoothed data was used for the ROI-to-ROI functional connectivity analyses and pattern similarity analyses. For the purpose of all functional connectivity analyses, additional preprocessing steps were applied. Outlier volumes were identified in functional images using the Artifact Detection Toolbox (ART) and a CompCor strategy was used for control of physiological and movement artifacts, both implemented in the CONN software, ver. 17.f. (Whitfield-Gabrieli & Nieto-Castanon, 2012).

### fMRI data analysis

fMRI data was analysed in line with three different approaches: univariate activation analysis, functional connectivity analysis and multivoxel pattern analysis (representational similarity analysis). The assumptions and rules of these approaches differ substantially, yet they were used to answer different questions and provide complementary information (Dimsdale-Zucker & Ranganath, 2019; Poldrack & Farah, 2015). A standard univariate fMRI analysis approach was used to examine the difference at each voxel between the averages across experimental conditions (emotion categories). Functional connectivity analysis was used to investigate if there was a temporal synchronization in the activation of different brain regions related to experimental conditions (emotion categories). Finally, representational similarity analysis (RSA) was used to examine the correlations between multivoxel activity patterns for each single trial (word pair) across encoding and recognition phases.

#### Univariate activation analysis

fMRI data was initially analyzed with SPM12 based on a mass-univariate approach and general linear models (GLMs). At the subject level, each single event was modeled with onset and duration corresponding to the presentation of a word pair, together with a response given afterwards (as it was not sufficiently separated in time from a word pair presentation). To account for movement-related variance, six nuisance regressors were included as representing the differential of the movement parameters from the realignment. Data was high-pass filtered (1/128 Hz) and convolved with a standard canonical hemodynamic response function (HRF) to approximate the expected blood-oxygen-level dependent (BOLD) signal. Because individual brain masks not always included all voxels comprising anatomically defined (based on the MNI coordinates) PRC region, we used the Imcalc tool (http://tools.robjellis.net/) for the SPM toolbox and constructed individual masks for the purpose of the analyses presented here. In order to incorporate all voxels comprising the PRC into individual masks, we used the logical “OR” expression and the masking threshold of –Inf. The resulting individual masks were included in the univariate analyses, as described below.

In the first analysis, the interaction between emotion and memory performance was examined for encoding and recognition sessions separately. Functional volumes from encoding session were split into conditions along the factors of emotion (DIS, FEA, NEU) and subsequent memory performance (corr, incorr). Thus, the following experimental conditions were specified: DIS corr, DIS incorr, FEA corr, FEA incorr, NEU corr, NEU incorr and miss (no response in subsequent recognition test). The number of trials falling into each condition was dependent on individual subsequent memory performance and varied between subjects (ranging from 10 to 35). Given that numerous subjects had less than 10 trials in the incorrect conditions, only correct conditions were further included in the group-level analysis. At the group level, flexible factorial design (Ashburner et al., 2014) was used because of the abovementioned variability in the number of trials among subjects and possible subject effects. Whole-brain random-effects contrasts were evaluated to obtain estimates of activity in response to each trial type relative to implicit baseline. Event-related stick-function regressors were used to perform ANOVA with emotion (three levels: DIS, FEA, NEU) as a within-subject factor, as well as a subject factor. The maps were created for the following directional T-contrasts: EMO – NEU (positive effect of emotion), DIS – FEA and FEA – DIS (difference between emotion categories).

Functional volumes from recognition session were split at the subject level into conditions according to emotion (DIS, FEA, NEU), memory performance (corr, incorr) and novelty (old, new). As a result, the following conditions were specified: old DIS corr, old DIS incorr, old FEA corr, old FEA incorr, old NEU corr, old NEU incorr, analogical conditions for new word pairs, and miss (no response). Once again, the number of trials in each condition was dependent on individual memory performance (ranging from 10 to 35 for the old and from 10 to 25 for new conditions). Given that numerous subjects had less than 10 trials in the incorrect conditions, again only correct conditions were further included in the group-level analysis. Similar to data analysis from encoding, at the group level, ANOVA was performed using flexible factorial design (Ashburner et al., 2014), with the following factors: emotion (three levels: DIS, FEA, NEU) and novelty (two levels: old, new) as within-subject factors, as well as a subject factor. Maps were created for the following directional T-contrasts: EMO – NEU (positive effect of emotion), DIS – FEA and FEA – DIS (difference between emotion categories). In order to test the positive effect of emotion on correct memory of old word pairs while controlling for more general emotional effects of new word pairs (Hofstetter, Achaibou, & Vuilleumier, 2012), maps were created for the following contrasts: old EMO – NEU masked exclusively (p < .05) by new EMO – NEU (positive effect of emotional memory), old DIS – FEA masked exclusively (p < .05) by new DIS – FEA, as well as old FEA – DIS masked exclusively (p < .05) by new FEA – DIS (difference between memorized emotions).

The second analysis was similar to the first one and also examined the interaction between emotion and memory performance during correct encoding and recognition. However, in order to isolate the effects of basic emotions (disgust, fear), modulation by the level of arousal was regressed out (Artur Marchewka et al., 2016), (A Marchewka et al., 2016). In order to perform this kind of operation, at the subject-level, individual arousal ratings collected during the third experimental session were included in the model. To this end, for each subject a vector representing subject’s individual ratings of arousal was convolved with HRF and added as an additional regressor of no interest.

Event-related stick-function regressors independent from the effects of arousal were subsequently used to perform ANOVAs at the group level in a manner described above.

The third analysis was performed to further examine how the general effects of memory were modulated by the individual ratings of each basic emotion (fear, disgust), affective dimensions (valence, arousal), and impact. At the subject-level, a vector representing subject’s individual ratings wa used as a linear (first-order) modulator (Artur Marchewka et al., 2016; M. Wierzba et al., 2018) of respective experimental conditions (corr and incorr at encoding; old corr, old incorr, new corr, new incorr at recognition). Conditions with zero variance in a parameter vector were discarded from further analysis. A single parameter was added to the model at a time to avoid orthogonalization effects between parameters (Mumford, Poline, & Poldrack, 2015). Furthermore, each parametric value was mean-centered to orthogonalize this variable with respect to the corresponding condition variable. Altogether, for each subject five additional models were created corresponding to the respective parameters: disgust, fear, valence, arousal, and impact. The effects (stick contrasts) of parametrically modulated conditions were then included in the group-level analyses. At the group level, ANOVAs were performed for each parameter separately, by defining a within-subject factor of memory performance (corr and incorr at encoding; old corr, old incorr, new corr, new incorr at recognition). The positive correlation was tested for each emotion parameter and each condition (the higher value of disgust, fear, arousal and impact parameter, the more emotional a given word pair). Only in the case of valence, negative correlation was analyzed (the lower value of valence parameter, the more emotional a given word pair).

If not stated otherwise, in all the analyses, a voxel-wise height threshold of p < .001 (uncorrected) combined with a cluster-level extent threshold of p < .05, corrected for multiple comparisons using the family-wise error (FWE) rate (Flandin & Friston, 2016) was applied in the whole brain analyses. The small volume correction (svc.) (Worsley et al., 1996) was applied using anatomical masks corresponding to the regions-of-interest (ROI) selected based on a priori hypotheses about the engagement of MTL in emotion and memory interactions. Within each of these ROIs, we considered activations whose effects survived the small volume FWE correction at the voxel level. This type of correction for multiple comparisons was previously used in works related to the topic of emotion and memory due to a small volume of subcortical structures of interest (Domínguez-borràs et al., 2017; Richter, Cooper, Bays, & Simons, 2016b). The coordinates of significant effects were reported in the Montreal Neurological Institute (MNI) space and labeled according to Automated Anatomical Labeling (AAL2) (Rolls, Joliot, & Tzourio-Mazoyer, 2015a) atlas with the use of bspmview (http://www.bobspunt.com/bspmview). Results were visualized with the use of MRIcroGL (http://www.mccauslandcenter.sc.edu/mricrogl/home).

#### Region-of-Interest localization and analyses

Based on the available literature, several regions of interest (ROI) were selected to further test specific hypotheses concerning differences in emotional modulation of successful encoding (followed by correct recognition). The ROI analysis was performed using a MarsBaR toolbox (Brett, Anton, Valabregue, & Poline, 2002). Contrast estimate values were extracted based on the subject-level SPM models used in the first univariate ANOVA analysis mentioned above in the following regions, bilaterally: amygdala (AMY), perirhinal cortex (PRC), hippocampus (HC) and parahippocampal gyrus (PHG). Anatomical masks for the ROI analysis were specified according to the AAL2 (Rolls, Joliot, & Tzourio-Mazoyer, 2015b) atlas implemented in the WFU PickAtlas (Maldjian, Laurienti, Kraft, & Burdette, 2003) toolbox, version 3.0.5. The PRC masks used in these analyses were manually segmented as described in (Ritchey et al., 2015).

As for emotion effects in these ROIs during encoding, first we performed an rm ANOVA on the contrast estimates extracted from selected regions, with the ROI (8 levels: AMY, PRC, HC, PHG, each left and right) and emotion (3 levels: disgust, fear, neutral) as within-subject factors. To unpack these effects, we run a post hoc analysis of simple effects restricted to the two emotion categories (DIS, FEA). Specifically, we performed a paired t-test for dependent samples to directly compare the mean contrast estimates between disgust- and fear-related stimuli all the ROIs.

As for emotion effects in these ROIs in the recognition data, we performed an rm ANOVA on the contrast estimates for old word pairs from selected regions, with the ROI (8 levels: AMY, PRC, HC, PHG, left and right) and emotion (3 levels: disgust, fear, neutral) as within-subject factors. Second, we performed a post hoc analysis of simple effects, namely a paired t-test for dependent samples to directly compare the mean contrast estimates between correctly recognized old disgusting and fearful stimuli in all the ROIs.

All the reported ANOVAs were performed with the Greenhouse-Geisser adjustment to the degrees of freedom when the assumption of sphericity was violated. Post hoc comparisons were performed using the Bonferroni adjustment for multiple comparisons.

#### fMRI functional connectivity analysis

Functional connectivity analysis aimed at investigating the temporal synchronization between the activation of brain regions involved in memory and emotion interactions was performed using the CONN software, ver. 17.f. (Whitfield-Gabrieli & Nieto-Castanon, 2012). Task-related connectivity changes were analyzed based on the subject-level SPM models from the first of univariate analyses described above. Experimental conditions with onsets and durations were imported from those models. Structural and functional images had been already preprocessed in SPM 12, including a normalization to MNI-space. An additional step of ART-based scrubbing was run for the outlier scans identification and a first-level covariate containing the outlier scans was created for each subject and session. The ROI-level analyses were performed using the unsmoothed data in order to avoid spillage from nearby regions (Whitfield-Gabrieli & Nieto-Castanon, 2017). The default ROI masks were imported for cortical and subcortical areas (AAL2 atlas), and a few commonly studied networks and areas as described above.

In a denoising step, linear regression was applied in order to remove unwanted motion, physiological, and other artifactual effects from the BOLD signal before computing connectivity measures. Three possible sources of confounds were defined: BOLD signal from the white matter and CSF masks, within-subject covariates (movement and scrubbing parameters) and the main effects of task conditions (direct BOLD signal changes associated with the presence/absence of a task) and regressed out. The high-pass filter of [0.008 Hz – Inf] was applied to keep higher-frequency information related to the task.

At the subject-level, generalized PsychoPhyisiological Interaction (gPPI) analysis (McLaren et al., 2012) was used to compute the interaction between the seed/source BOLD timeseries and a chosen condition-specific factor when predicting each voxel or target ROI BOLD timeseries. Bivariate regression was computed for each pair of source and target ROIs (ROI-to-ROI analysis). The obtained regression coefficients were used for the group-level analysis of increases and decreases in functional connectivity. Similar to the univariate activation analyses, the following contrasts were tested for encoding and recognition sessions: EMO – NEU, DIS – FEA, and FEA – DIS. The results were reported at a two-sided threshold of p < 0.05, with false discovery rate (FDR) (Benjamini & Hochberg, 1995) and correction for multiple comparisons at the seed-level.

#### fMRI multivariate pattern analysis

Previous analyses of brain activity were performed on the encoding and recognition data separately. However, another very interesting aspect is how emotion modulates the relationship between brain activation during encoding and retrieval (Ritchey, Wing, LaBar, & Cabeza, 2013) (here more specifically: recognition). In order to examine emotion modulation of the similarity of transient, stimulus-evoked BOLD activity between encoding and recognition, we performed a split-half correlation analysis within the framework of representational similarity analysis (RSA) (Kriegeskorte, 2008). In this multivariate approach, data from individual voxels within a region of interest are jointly analyzed, which corresponds to a contemporary view on the brain representation of different mental states (Kragel & LaBar, 2014). A general rule of a split-half correlation analysis is that the mean correlation for the non-matching trial pairs is subtracted from a mean correlation for the matching trial pairs. It is assumed that if an area represents certain trial-specific information in a distributed manner (over voxels), then correlations (over voxels) of matching trials may be higher than correlations of non-matching trail pairs.

The first step of this analysis was creating the separate subject-level GLM models for each of 105 encoding trials and each of 180 recognition trials to estimate the single-trial response. A Least-Square Single (LSS) method was used to model each trial as a regressor of interest and combine the remaining trials into a single nuisance regressor (Mumford, Turner, Ashby, & Poldrack, 2012). Each trial’s onsets and durations were extracted from the existing subject-level models as described in the first univariate analysis above. The obtained single-trial beta images were then sorted identically for encoding and recognition, and concatenated into 4D NIfTI format for further analyses.

Then, the split-half correlation RSA was performed with the CoSMO MVPA (Oosterhof, Connolly, & Haxby, 2016) toolbox and in-house MATLAB scripts. Both searchlight (Kriegeskorte et al., 2006) and ROI-based methods were used. First, we applied a searchlight method, where a sphere-shaped mask ‘travels’ through the brain, and at each location a measure of interest (here: a correlation coefficient) is assigned to the center voxel of the sphere, resulting in a whole-brain map (here: of correlation differences). In order to generate searchlight maps for each participant, a sphere with a radius of 3-voxels (on average, 110 voxels per sphere) was defined as a mask. In a given searchlight sphere, the multivoxel response patterns for each encoding and recognition trial (item) were extracted from beta images. Next, for each trial the similarity between encoding and recognition response patterns was estimated using a pair-wise Pearson correlation. Finally, the resulting correlation coefficients were transformed using Fisher’s r-to-z formula. In total, multiple (over 11000) correlation coefficients were assigned to each voxel, 105 of which represented the voxelwise correlation between matching encoding-recognition trial pairs, whereas the remaining values represented the voxelwise correlation between the non-matching encoding – recognition trial pairs.

Second, the data was prepared for the RSA analysis applying the ROI-beased method. We expected that the similarity of brain activation patterns between encoding and recognition could arise not only from the reactivation of perceptual processes, but also from higher-order cognitive and affective processes in MTL regions. Therefore, all the ROIs selected for the ROI univariate analysis were also selected for the split-half correlation RSA. Similarly to the searchlight approach just described, for each trial the similarity between encoding and recognition response patterns in each ROI was estimated using a pair-wise Pearson correlation. The resulting Fisher-transformed r-to-z score was assigned to each ROI. In total, multiple correlation coefficients were assigned to each ROI, representing either matching or non-matching encoding – recognition trials.

Finally, the index of encoding-retrieval similarity (ERS) (Ritchey, Wing, Labar, et al., 2013; Tulving & Thomson, 1973) was calculated for each sphere and ROI per participant, as a mean correlation for the matching pairs minus a mean correlation for the non-matching trial pairs (one of the possible forms of split-half correlation analysis). To this end, positive values were put on the diagonal of the contrast matrix, and small negative values were put on the off-diagonal area of the contrast matrix, which is depicted in Fig. 6. The resulting single value was assigned to the central voxel of each sphere and each ROI. To further explore ERS corresponding to different experimental conditions, a contrast matrix was created for the resulting values of ERS index to directly compare different levels of correctness (2 levels: correct, incorrect) and emotion (3 levels: disgust, fear, neutral) factors. In the case of searchlight method, the t values resulting from the contrasts were again assigned to each central voxel of the sphere, and the obtained maps of t-values were tested against a null hypothesis of zero using one-sample t-test across subjects. Additionally, one sample t-test across subjects was also performed on the single t-values assigned to each ROI.

**Fig. 6.**
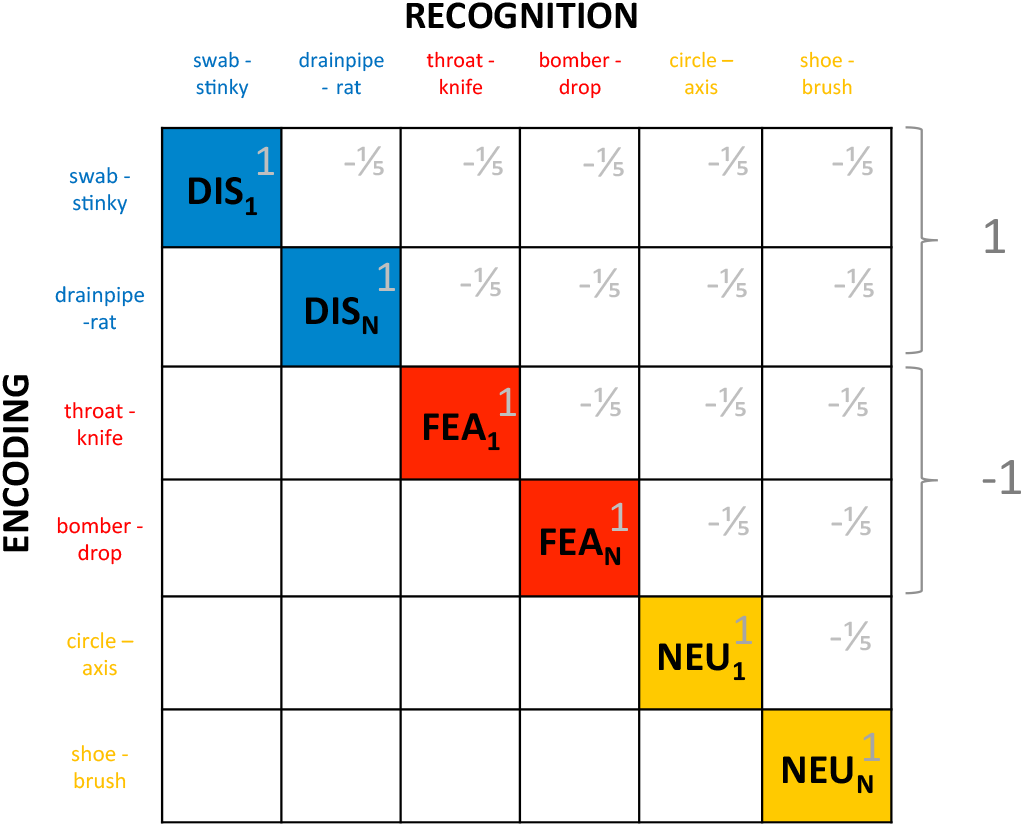
Schematic of a contrast matrix of the ERS calculated for the matching (diagonal) and non-matching (off-diagonal) pairs of encoding and retrieval trials, across emotion categories. DIS - disgust, FEA - fear, NEU - neutral. The contrast matrix was also modified to directly compare the ERS values between emotion and correctness conditions.

#### Encoding – reinstatement analysis

Due to variability in the ERS index across trials, the correspondence between encoding and recognition might not be fully illustrated by the results of multivariate ERS analysis averaged for each across trials for each emotion category. Thus, the goal of the final analyses (based on (Danker et al., 2017)) was twofold. First, we aimed at investigating whether the recognition brain activity could be predicted by encoding activity in regions related to emotion and memory interactions. Specifically, we queried the retrieval data for voxels, whose trial-by-trial activity was predicted by (correlated with) trial-by-trial encoding activity averaged within each regions of interest (ROIs): bilateral AMY, HC, PHG and PRC. Second, we wanted to investigate the relationship between the encoding activity and subsequent reinstatement, as measured with the ERS index. Thus, we queried the encoding data for voxels, whose trial-by-trial activity correlated with trial-by-trial ERS calculated for emotion-specific masks. These masks were created from the following contrasts: DIS – FEA, FEA – DIS and NEU – (DIS + FEA), including voxels at p < .001 uncorrected.

In the first reinstatement analysis, we initially used in-house scripts based on Marsbar to extract contrast estimates in the abovementioned ROIs for each encoding trial. As presented in Fig. 7, these trial-by-trial contrast estimates were further used as parametric regressors applied to the recognition data in a model that accounted for parametric effects within each emotion and correctness condition. In other words, each old trial at recognition obtained a parametric value corresponding to the AMY, PRC, HC and PHG activation for that trial at encoding. Each individual subject’s model included the following regressors for old conditions: DIS corr, DIS corr parametric, DIS incorr, DIS incorr parametric, FEA corr, FEA corr parametric, FEA incorr, FEA incorr parametric, NEU corr, NEU corr parametric, NEU incorr, NEU incorr parametric; and new DIS corr, new DIS incorr, new FEA corr, new FEA incorr, new NEU corr, new NEU incorr and miss. This approach was aimed to identify regions that exhibit trial-by-trial retrieval activity correlated with trial-by-trial AMY, HC, PHG and PRC encoding activity across experimental conditions.

**Fig. 7.**
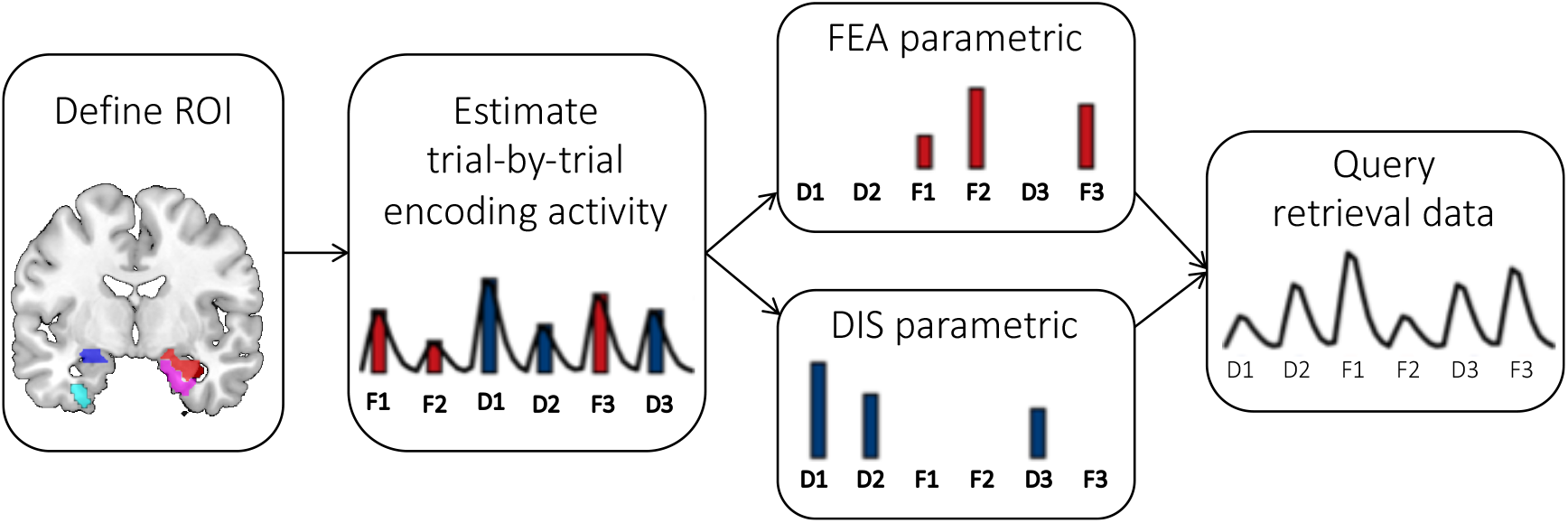
Schematic of the encoding – reinstatement analysis with parametric modulation of recognition data by trial-specific ROI activation during encoding. D1 – 1^st^ disgust trial, D2 – 2^nd^ disgust trial, D3 – 3^rd^ disgust trial, F1 – 1^st^ fear trial, F2 – 2^nd^ fear trial, F3 – 3^rd^ fear trial (modified after (Danker et al., 2017)).

In the second reinstatement analysis, emotion-specific masks from encoding were used as ROIs for the ERS analysis as described above (DIS – FEA used for disgust-related trials, FEA – DIS used for fear-related trials and NEU – (DIS + FEA) used for neutral trials). These trial-by-trial ERS estimates were used as parametric regressors applied to encoding data in two models, one of which accounted for parametric effects of ERS within correctness factor only. Thus, as presented in Fig. 8, each trial at encoding obtained a parametric value corresponding to the ERS value for that trial. In this model, ERS values for disgust-related, fear-related and neutral trials were concatenated into a single parametric regressor and this analysis was meant to find encoding regions that predicted trial-by-trial ERS across emotions. Since ERS values for each emotion were calculated based on different trials and different set of voxels, they were separately mean-centered. Each individual subject’s model included the following regressors: corr, corr parametric, incorr, incorr parametric and miss. Next, in order to investigate the various effects for emotions, another parametric model was constructed which included separate regressors for each level of emotion and correctness factors. Here, each individual subject’s model included the following regressors: DIS corr, DIS incorr, FEA corr, FEA incorr, NEU corr, NEU incorr and miss. In general, using this approach, we could identify regions exhibiting activation during encoding that correlated with broad similarity in patterns of activity between encoding and their corresponding retrieval trials, across experimental conditions.

**Fig. 8.**
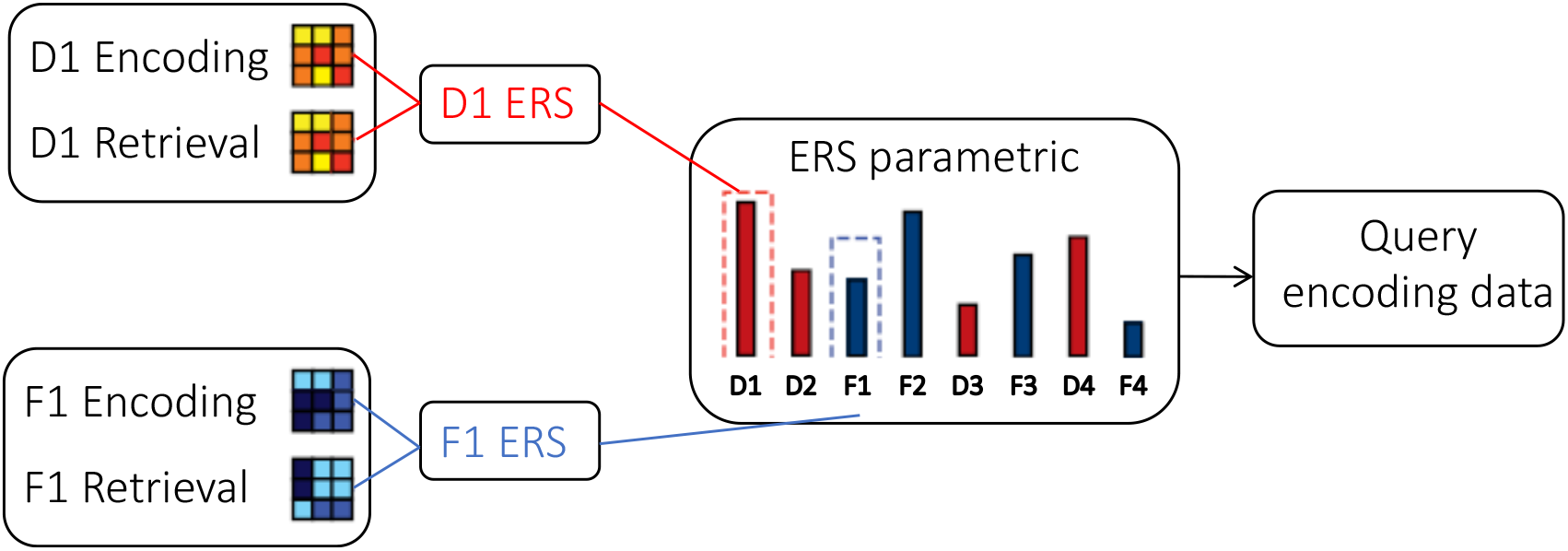
Schematic representation of the encoding – reinstatement analysis with parametric modulation of encoding data by the trial-by-trial ERS index. D1 – 1^st^ disgust trial, D2 – 2^nd^ disgust trial, D3 – 3^rd^ disgust trial, D4 – 4^th^ disgust trial, F1 – 1^st^ fear trial, F2 – 2^nd^ fear trial, F3 – 3^rd^ fear trial, F4 – 4^th^ fear trial (modified after (Danker et al., 2017)).

## Acknowledgements

We would like to thank: Dawid Droździel and Bartosz Kossowski for their technical help with the MR scanner, Michał Szczepanik for his help to program the experiment, Karolina Golec for her help with data collection, James Keidel for his advice on the RSA analyses, Lila Davachi for her inspiring advice on encoding - reinstatement analysis, Oded Bein for his useful comments on the manuscript. This work was funded by the National Science Centre Poland (2012/07/B/HS6/02112 to AG, 2015/19/N/HS6/02376 and 2016/20/T/HS6/00600 to MR) and the Foundation for Polish Science (START 071.2017 to MR).

