## Supplementary materials for "Distinct medial-temporal lobe mechanisms of encoding and amygdala-mediated memory reinstatement for disgust and fear"

### Supplementary analyses and figures

#### Recognition rate

The analysis of recognition rate (proportion of correct responses) revealed that the participants remembered both categories of emotional word pairs better than neutral word pairs and that disgust was remembered better than fear (main effect of emotion [F(2,102) = 19.78, p < .001, η2 = .28]). More specifically, this relationship differed depending on the novelty of word pairs (interaction effect between emotion and novelty [F(2,102) = 19.48, p < .001, , $\eta^2$ = .28]).

Among old word pairs, the recognition rate was higher for disgust than fear, as well as higher for disgust than neutral, and higher for fear than neutral (p < .001) pairs. Among new word pairs, it was only lower for fear than neutral pairs. Most importantly, however, recognition rate was higher for old than new word pairs related to disgust, it did not significantly differ (although in the same direction) between old and new word pairs related to fear, but it was lower for old than new (p = .008) neutral word pairs. In other words, we observed that emotionality of word pairs differently influenced the recognition of old and new word pairs.

Although the participants were better in recognizing old than new word pairs related to disgust and fear, in the case of neutral word pairs they gave more correct responses when recognizing new word pairs. When we excluded the effects related to neutral word pairs and compared only the effects of basic emotion categories, we could still find significant differences between emotion categories and a main effect of emotion [F(1,51) = 22.14, p < .001, $\eta^2$ = .05], but neither an effect of novelty, nor the interaction between emotion and novelty. This analysis showed simply that word pairs related to disgust were recognized better than word pairs related to fear (p < .001), regardless of their novelty.

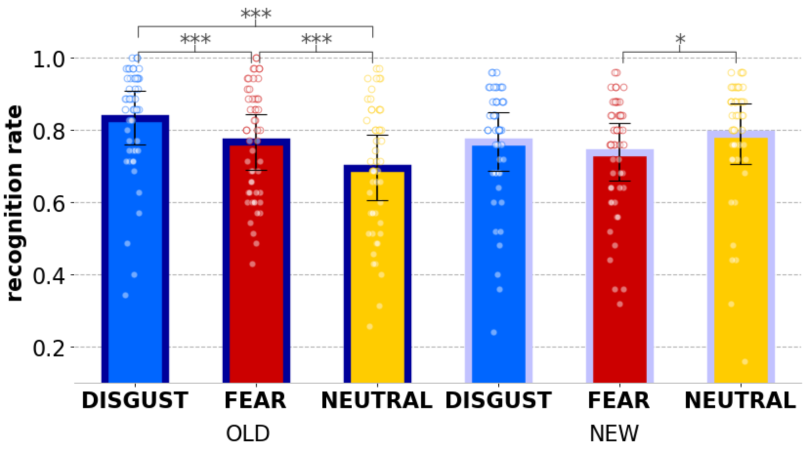


Fig. S1 Recognition rate (proportion of correct responses) with emotion and novelty factors; error bars represent one standard deviation, dots represent individual subjects' scores; *** p < .001, * p < .005.

In order to isolate the effects of different emotion categories (disgust, fear), we aimed at controlling for the level of arousal evoked by each word pair. Therefore, we included the difference in the level of arousal ratings between disgust- and fear-related word pairs as a covariate in the analysis of recognition rate. Both old (main effect of emotion [F(1,50) = 13.80, p < .001, $\eta^2$ = .22]), and new [F(1,50) = 4.32, p = .043, $\eta^2$ = .08] word pairs evoking disgust were still better recognized than word pairs evoking fear, and there was no interaction effect between emotion and a covariate of arousal difference.

#### Semantic congruency

An additional one-way ANOVA with one factor of emotion (3 levels: disgust, fear, neutral) was performed for the final verification of experimental manipulation and to indicate if there were no differences in mean semantic congruency of all word pairs used in the experiment (n = 180). It revealed no significant effects [$F(2,177) = .561$, $p = .572$], as depicted in Fig. S2.

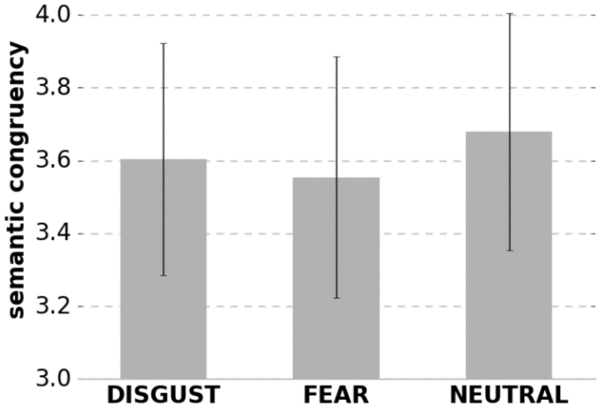


Fig. S2 Results of one-way ANOVA on the mean ratings of semantic congruency between words forming word pairs with emotion factor. Error bars represent standard deviation.

#### Brain mechanisms – encoding

##### Univariate whole brain analysis, model without regressing out arousal

We found that successful encoding of emotional items engaged extensive regions of activity, including prefrontal, temporal and posterior regions, as well as bilateral amygdala (AMY). However, a direct comparison of emotions showed that successful encoding of disgust and fear engaged distinct brain regions. Specifically, disgust (DIS > FEA) engaged left dmPFC, middle frontal gyrus (MFG), bilateral MTG, left OCC, left ANG, as well as left AMY and PRC, whereas fear (FEA > DIS), engaged bilateral MTG, left middle OCC, right PCUN, left PHG and right HC.

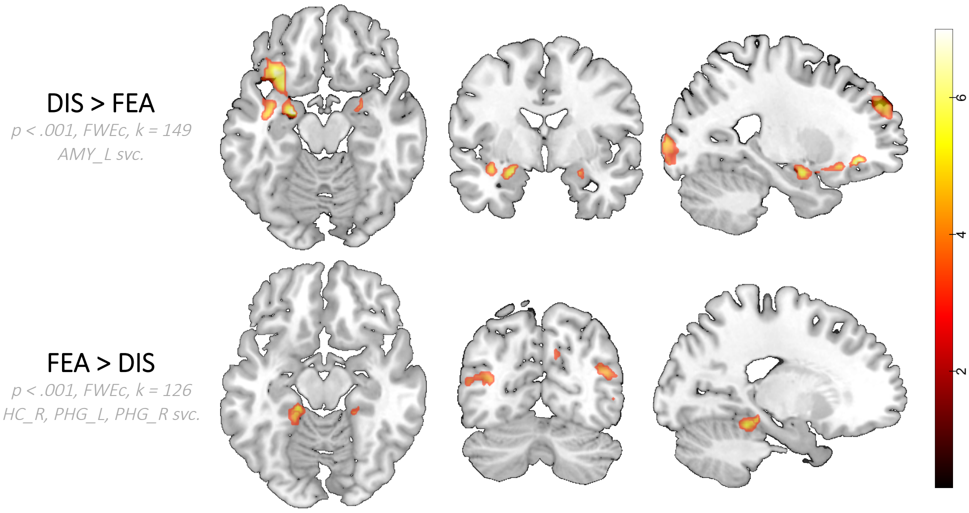


Fig. S3 Brain activation differences between basic emotion conditions (disgust and fear), during successful encoding, in both directions: DIS > FEA corr – disgust > fear correct, FEA >

DIS corr – fear > disgust correct; FWEc – cluster-level FWE–corrected; k – cluster extent, svc. – small volume correction; colour scale represents a range of t-values.

#### Modulation by affective parameters

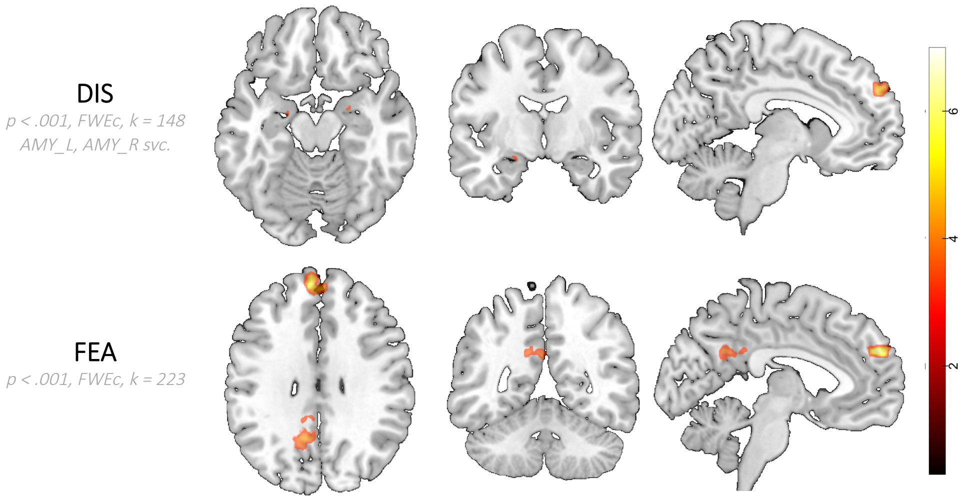


Fig. S4 Brain activation during encoding modulated by affective parameters of disgust and fear (positive effects); FWEc – cluster-level FWE–corrected; k – cluster extent, svc. – small volume correction; colour scale represents a range of t-values.

#### Brain mechanisms – recognition

##### Univariate whole brain analysis, model without regressing out arousal

We found that during correct recognition, emotional conditions (old EMO > NEU masked exclusively by new EMO > NEU) were related to the activations only in left hemisphere, including prefrontal, posterior regions and AMY. Also disgust as compared to neutral (old DIS > NEU masked exclusively by new DIS > NEU) engaged left AMY, whereas fear compared to neutral (old FEA > NEU masked exclusively by new FEA > NEU) engaged bilateral MTG, left PCC, right CS, right LG, left dmPFC, and right HC.

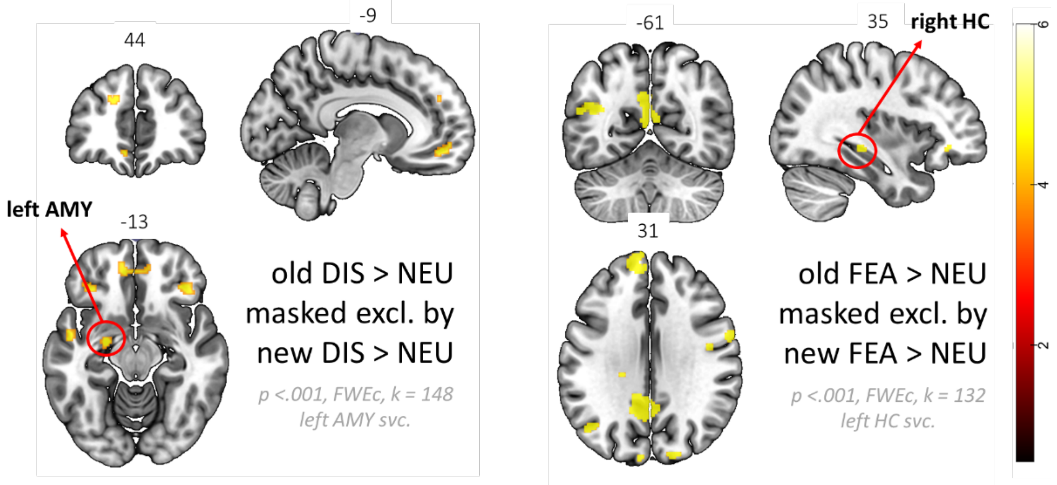

Fig. S5 Differences in brain activation during successful recognition between basic emotion conditions (disgust, fear) and neutral condition, in both directions. Effects in old word pairs were exclusively masked for the same effects in new word pairs. DIS > NEU – disgust vs. fear correct, FEA > NEU – fear vs. disgust correct; FWEc – cluster-level FWE–corrected; k – cluster extent, svc. – small volume correction; colour bar represents a scale of t-values; xyz coordinates in MNI space given above each brain slice.

However, a direct comparison of disgust and fear did not reveal the differences similar to those found during encoding, namely correct recognition of disgust (old DIS > FEA masked exclusively by new DIS > FEA) revealed activations in left MFG, IFG and IPL as well as marginally in the left PRC, whereas correct recognition fear (old FEA > DIS masked exclusively by new FEA > DIS) showed activations in right superior parietal lobule (SPL) and postcentral gyrus.

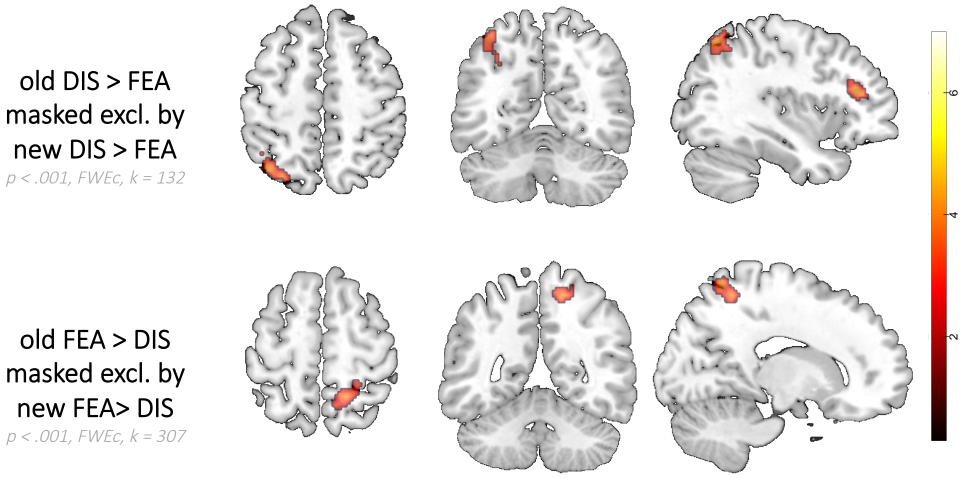


Fig. S6 Differences in brain activation during successful recognition between basic emotion conditions (disgust and fear), in both directions. Effects in old word pairs were exclusively masked for the same effects in new word pairs. DIS > FEA corr – disgust vs. fear correct, FEA > DIS corr – fear vs. disgust correct; FWEc – cluster-level FWE–corrected; k – cluster extent; colour scale represents a range of t-values.

#### Modulation by affective parameters

An additional analysis showed that during recognition, brain activity was modulated by individual differences in experienced emotion (as operationalized by subjective affective ratings) of disgust in bilateral MTG, left dmPFC, IFG, INS, PCC and right PCUN, whereas modulation by fear was observed only in left supramarginal gyrus (Supplementary Fig. 9). No regions related to memory or emotion processing were found to be modulated by other affective parameters: impact, valence, arousal.

FEA
*p < .001, FWEc, k = 307*

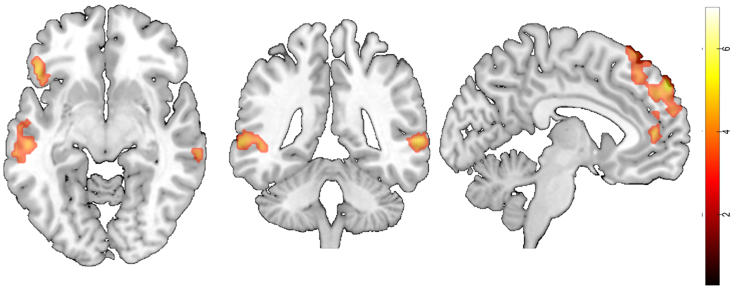

Fig. S7 Brain activation during recognition modulated by affective parameters of disgust and fear (positive effects); FWEc – cluster-level FWE–corrected; k – cluster extent; colour scale represents a range of t-values.

#### Supplementary tables

Table S1. List of all the word pairs created for the experiment from NAWL words and used in the encoding session. CAB presentation order; 1 – disgust, 2 – fear, 3 – neutral.

| Word pair | Emotion category |
|---|---|
| mydliny - czyścić | 1 |
| symptom - chory | 2 |
| gąsienica - larwa | 1 |
| gaz - wybuch | 2 |
| karafka - porto | 3 |
| byk - uciekać | 2 |
| nos - palec | 3 |
| twierdza - wampir | 2 |
| gardło - nóż | 2 |
| słowo - łaciński | 3 |
| bombowiec - zrzut | 2 |
| czopek - dupa | 1 |
| rekin - paniczny | 2 |
| błazen - służyć | 1 |
| nożyczki - nacinać | 2 |
| kura - bezgłowy | 1 |
| zajazd - przyjmować | 3 |
| przecinek - zdanie | 3 |
| bażant - zając | 3 |
| kula - dziura | 2 |
| szmata - ohydny | 1 |
| skrzynia - przenosić | 3 |
| członek - organ | 1 |
| chwast - oset | 1 |
| kartoteka - lustrować | 2 |

| | |
|---|---|
| nieruchomy - wynosić | 2 |
| silnikowy - stać | 3 |
| stempel - kopia | 3 |
| kamera - szpiegować | 2 |
| pień - czeremcha | 3 |
| kosa - plaster | 2 |
| dentysta - ząb | 2 |
| talerz - podawać | 3 |
| mucha - padlina | 1 |
| język - opisywać | 3 |
| napad - rewolwer | 2 |
| patelnia - łyżka | 3 |
| dzbanek - kubek | 3 |
| ścieki - oczyszczać | 1 |
| wzrost - metr | 3 |
| kierowca - radar | 2 |
| zastrzyk - zemdleć | 2 |
| wstręt - gorzki | 1 |
| punkt - atramentowy | 3 |
| broń - policyjny | 2 |
| krzesło - siadać | 3 |
| szafa - bluzka | 3 |
| brodawka - maszkara | 1 |
| talar - miedziany | 3 |
| gówno - gołąb | 1 |
| urzędnik - biurowy | 1 |
| pirania - przerażający | 2 |
| slogan - tendencyjny | 1 |
| koza - poroże | 1 |
| mafia - zlecać | 2 |
| dynia - marchwiowy | 3 |
| żaba - odrażający | 1 |
| nadzorca - kurator | 2 |
| piwnica - krata | 2 |
| adresat - znaczek | 3 |
| anonim - podstępny | 2 |
| łazienka - grzebień | 3 |
| leksykon - nazywać | 3 |
| kartka - pędzel | 3 |
| kanał - szczur | 1 |
| jama - lis | 2 |
| świętokradztwo - haniebny | 1 |
| cecha - profil | 3 |
| toaleta - karaluch | 1 |
| parasol - suszyć | 3 |

| | |
|---|---|
| zebra - gromadny | 3 |
| świnia - szczecina | 1 |
| moździerz - rakieta | 2 |
| bagno - zmurszały | 1 |
| Koran - maniakalny | 2 |
| but - szczotkować | 3 |
| ksiądz - radykał | 2 |
| powieka - oko | 3 |
| dokument - sprawozdanie | 2 |
| kopalnia - przemysłowy | 2 |
| tampon - wilgotny | 1 |
| buk - kora | 3 |
| amunicja - sabotaż | 2 |
| zgnilizna - obrzydliwy | 1 |
| robak - gnić | 1 |
| przedstawiciel - senat | 1 |
| termin - zabiegany | 2 |
| lupa - mniejszy | 3 |
| regał - segregator | 3 |
| piła - ostrze | 2 |
| owad - chrząszcz | 1 |
| prześcieradło - kłaść | 3 |
| koniak - skacowany | 1 |
| cygaro - cuchnący | 1 |
| topór - brutalny | 2 |
| wódka - kiełbasa | 1 |
| magnes - matowy | 3 |
| kurtka - szal | 3 |
| kosz - odpady | 1 |
| stołówka - obskurny | 1 |
| mdłości - spleśniały | 1 |
| komisja - wniosek | 2 |
| mur - przyciskać | 2 |
| żółć - wymiotować | 1 |
| awersja - tytoniowy | 1 |
